# RNA Organelles in DNA-based Artificial Cells Provide Spatial Aptamer Functions and Enhanced Signal Processing

**DOI:** 10.64898/2026.06.22.733869

**Authors:** Maria Vonk-de Roy, Daniel Hoenders, Laia Civit, Julián Valero, Andreas Walther

## Abstract

Naturally occurring biomolecular condensates orchestrate key cellular processes by creating spatially distinct reaction environments, yet engineering synthetic condensates that combine structural programmability with spatioselectively encoded function remains challenging. Here we report multiphase DNA–RNA artificial cells (ACs) that embed functional RNA condensates as organelle-like compartments within programmable DNA core–shell ACs. A single thermal assembly protocol yields three-phase ACs comprising a glassy RNA core organelle embedded in a liquid-like DNA compartment, surrounded by a crosslinked DNA shell. The RNA organelles contain aptamer function, enabling selective protein recruitment and small-molecule activation, while the DNA scaffold provides independent addressability, regulates RNA-condensate size and enhances resistance to serum-mediated degradation. We further show that RNA chemistry can be used to adjust environmental responsiveness: unmodified RNA organelles undergo rapid degradation in serum and release captured protein cargo, whereas 2′-fluoro-modified RNA organelles remain stable for at least 24 h. Finally, by coupling transcriptional modules localized in the DNA core to cell-free protein translation in the surrounding medium, we establish sender–receiver communication between AC populations and self-actuating signal processing within individual DNA–RNA ACs. These results establish hybrid nucleic-acid ACs as programmable, spatially organized systems that couple compartment architecture, RNA molecular recognition and biochemical communication.

**TOC Figure:** 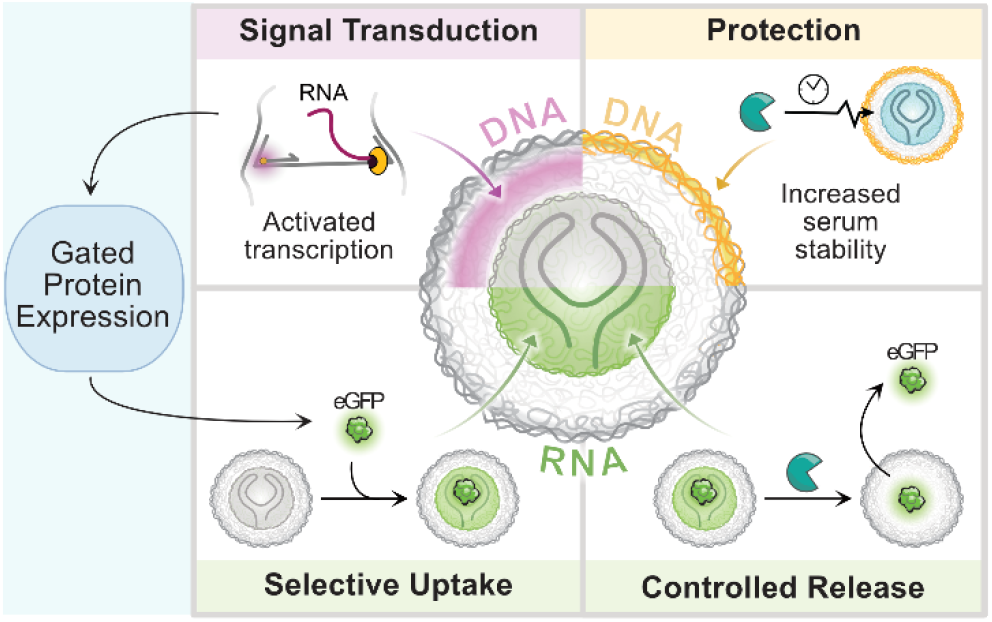

Multiphase DNA–RNA artificial cells integrate a protective DNA shell, a transcriptionally active DNA core, and a functional RNA organelle. Spatial compartmentalization enables signal generation, external protein expression, and selective recapture via RNA aptamers.

## Introduction

Membrane-free biomolecular condensates are central regulators of intracellular biochemistry, concentrating proteins and nucleic acids into dynamic compartments that control processes such as RNA metabolism, ribosome biogenesis, and signal transduction.^1–3^ These assemblies are formed via phase separation and enable cells to locally enrich reactants, which in turn spatially control reaction environments to coordinate complex biochemical pathways.^4^ Such condensates do not only appear in the cytoplasm, but RNA-rich condensates can also form within the DNA-rich cell nucleus.^5^

Inspired by these functions, synthetic condensate systems have been explored to create compartmentalized biochemical environments *in vitro*.^6^ Nucleic acids are especially attractive building blocks due to their sequence-inherent programmability. This programmability enables assembly through complementary base-pair interactions,^7–12^ or sequence-specific temperature-induced condensation mediated by multivalent ions.^6,13–17^ Present synthetic nucleic acid condensates for the bionanoscience field typically focus on DNA-based structures that have given rise to the field of all-DNA based artificial cells (ACs).^18^ In contrast, RNA – due to its additional hydroxy group and ability to fold into more complex secondary structures – is highly relevant for extending functionality inside nucleic acid-based ACs as it can encode a diversity of catalytic ribozyme actions, selective molecular recognition via aptamers and can be a regulator for protein translation.^19,20^ The integration of such functional RNA elements into phase-separated condensates in a controlled and spatially organized manner, however, remains challenging, with limited examples of functionalizing DNA nanostar condensates with aptamers^21^ or initiating transcription.^11^ We recently demonstrated in situ RNA condensate formation with DNA ACs formed by phase separation.^22^

Given that both DNA and RNA can undergo temperature-induced phase separation^7–12^, we raised the question whether a combined DNA–RNA system would allow for an integrated programmable structure formation with biochemical functionality within this hybrid condensate system, essentially recapitulating minimalistic aspects of the spatial organization found in living cells. Such multiphase architectures with distinct functional regions would enable controlled partitioning of biomolecules, localized reaction environments, and communication between compartments. These features are essential for constructing higher-order systems for synthetic biology and bionanoscience.

Here, we engineer DNA–RNA artificial cells (DR-ACs) that combine a programmable DNA condensate scaffold with functional RNA condensates in a unified multiphase architecture. We demonstrate that RNA condensates can be embedded as a distinct RNA “organelle” within a DNA core–shell condensate using a one-step thermal protocol, thereby establishing spatially separated domains with distinct functional roles. The RNA condensate phase retains aptamer-encoded molecular activity, while the surrounding DNA condensate remains independently addressable. Furthermore, the DNA scaffold modulates RNA organelle size and enhances resistance to serum-mediated degradation, which can be further enhanced by chemically modified RNAs, collectively enabling increased stability under biologically relevant conditions. Finally, coupling localized transcription within a DNA compartment to an external cell-free translation system enables protein production and selective capture within an RNA-protein aptamer organelle. This yields sender-receiver communication between ACs as well as integrated signal processing within single DR-ACs. Together, this approach establishes a platform for engineering spatially organized, functionally integrated DNA-RNA systems that bridge synthetic condensates and cellular organization.

## Results and Discussion

### Synthesis and optimization of multiphase DNA–RNA ACs

We engineer multiphase DR-ACs by extending our previously reported all-DNA-AC platform, in which poly(A_20_-x)_n_ and poly(T_20_-y)_n_ ssDNA polymers assemble into core–shell condensates using a rapid temperature ramp in the presence of 50 mM MgCl_2_.^6,13,15–17^ A_20_ and T_20_ are homo-DNA repeats of adenine and thymine, respectively, whereas x and y are defined nucleic acid barcodes. During heating, the adenine-rich poly(A_20_-x)_n_ undergoes liquid–liquid phase separation (LLPS) to generate micron-scale droplets, onto which the poly(T_20_-y)_n_ hybridizes through T_20_/A_20_ duplex formation during cooling, generating an interfacially crosslinked shell and thus a final core–shell AC architecture (**Error! Reference source not found.a**). While the A_20_ and T_20_ repeating units define the AC architecture, the barcodes x and y are programmable and enable post-assembly hybridization of fluorescent labeled oligonucleotides (Error! Reference source not found.**b**) for confocal laser scanning microscopy (CLSM) or other functionalizations. Error! Reference source not found.**c** shows an example of fluorescently labeled DNA-ACs generated from poly(A_20_-x)_n_ and poly(T_20_-y)_n_ (3:1 ratio).

To explore the incorporation of RNA modules into these DNA-ACs, we first investigate whether an eGFP binding RNA aptamer (AP3-RNA) undergoes LLPS under the same buffer, salt and temperature conditions.^23^ Indeed, comparable micron-scale condensates are formed, which are presumably held together via percolated phase-separated structures and potentially some RNA–RNA bonds.^6,14^ RNA condensates form robustly from HPLC-purified AP3-RNA (**Error! Reference source not found.d**), as well as when used directly from *in vitro* transcription mixtures (**Error! Reference source not found.e**), indicating that reaction components of the transcription system (including T7 RNA polymerase) do not measurably influence condensate formation under these conditions. Moreover, incorporation of Cy5-labeled UTP (UTP-Cy5) during transcription yields fluorescently labeled RNA condensates, enabling direct visualization by CLSM without additional staining (Error! Reference source not found.**f**).

Interestingly, combining AP3-RNA with poly(A_20_-x)_n_ and poly(T_20_-y)_n_ (16:3:1 ratio; 750:144:48 ng µL^-1^), and subjecting it to the same temperature ramp at 50 mM MgCl_2_, yields hybrid DR-ACs. These display a distinct three-phase architecture comprising an RNA-rich interior (termed “RNA organelle”) enclosed in a DNA-rich poly(A_20_-x)_n_ compartment and an outer poly(T_20_-y)_n_ DNA shell (Error! Reference source not found.**g,h)**. Because AP3-RNA exhibits no significant sequence complementarity to the x and y barcode regions (**Figure S1)**, we suggest that structure formation follows interfacial-tension consideration and nucleated processes. This implies that RNA condensates have a higher interfacial tension to the solution and the highest propensity for phase separation, whereas DNA parts have a lower one and thus assemble on the surface of RNA condensates.^24^ Consistent with this hypothesis, temperature ramps of AP3-RNA with either poly(A_20_-x)_n_ or poly(T_20_- y)_n_ alone produce concentric DR-ACs in both cases (**Figure S2**). Notably, poly(T_20_-y)_n_ forms a continuous, conformal layer around the RNA condensates, whereas poly(A_20_-x)_n_ yields a slightly discontinuous one. This suggests that both ssDNA polymers can interfacially bond to RNA condensate through non-specific interactions, for instance, via Mg^2+^ induced percolation at high temperatures.

Initial observations indicate that RNA organelles in DR-ACs are smaller and more monodisperse than the RNA condensates formed in the absence of DNA (Error! Reference source not found.**f–h**). To study the effect of DNA on RNA condensate size and distribution, we subjected seven AP3-RNA– poly(T_20_-y)_n_ samples with identical AP3-RNA concentrations (750 ng µL^-1^) at varying poly(T_20_-y)_n_ concentrations (ranging between 0 to 50 ng µL^-1^) to temperature-induced phase separation. Poly(A_20_- x)_n_ was omitted because poly(T_20_-y)_n_ is capable of forming well-defined shells by itself. The resulting RNA organelles indeed have a smaller diameter at elevated DNA concentrations (diameter: 2.0 µm, standard deviation: 0.8 µm) at 50 ng µL^-1^ poly(T_20_-y)_n_, whereas RNA condensates prepared in the absence of DNA are larger with a larger size distribution (diameter: 2.6 µm, standard deviation: 1.5 µm; **Figure S3**). Notably, poly(T_20_-y)_n_ concentrations below 12.5 ng µL^-1^ yield DR-ACs with patchy DNA shells, with clearly observable clustering behavior. To further tune RNA organelle dimensions in AP3-RNA–poly(T_20_-y)_n_ DR-ACs, we prepared samples containing 750 ng µL^-1^ AP3-RNA and 25 ng µL^-1^ poly(T_20_-y)_n_ and subjected them to temperature ramps with 11 different high temperature plateau times (15 – 300 s at 95 °C). Longer plateau times systematically increase condensate size and reduce particle number (**Figure S4**). This anti-correlation suggests that prolonged high-temperature incubation does not increase the fraction of phase-separated RNA, but instead promotes condensate coalescence, yielding fewer but larger particles.

Finally, we assessed whether RNA organelle formation within DR-ACs is a general phenomenon or limited to specific RNA sequences such as AP3-RNA. To this end, we tested 8 different RNA molecules, with varying length, sequence, and folding, including aptamers, random RNAs, and mRNA (34-1484 nt) to study their ability to undergo condensation in the absence of DNA, and to form DR-ACs with poly(A_20_-x)_n_ and poly(T_20_-y)_n_ (10:3:1 ratio; 500:144:48 ng µL^-1^; **Figure S5**). While some RNA sequences form weak or undetectable condensates on their own, all tested RNAs do incorporate in DR-ACs, albeit at reduced RNA enrichment or altered spatial distributions of compartments in certain cases. This demonstrates that the approach is broadly compatible with diverse RNA sequences, with a slight preference for a more robust co-condensation for longer RNAs. Since these experiments were performed under conditions optimized for AP3-RNA, further optimization of salt conditions may enhance incorporation efficiency for other short RNA sequences. Together, these findings show that RNA phase separation is compatible with DNA-AC assembly, enabling DR-AC formation with tunable RNA organelle size and incorporation of diverse RNA sequences.

### Structural and dynamic characterization of DR-ACs

We next investigate how decreasing the RNA concentration influences the internal organization of DR-ACs. To this end, we lowered the AP3-RNA concentrations from 750 to 375 and 188 ng µL^-1^, while keeping the poly(A_20_-x)_n_ and poly(T_20_-y)_n_ concentrations constant (**Figure 2a**). While the highest AP3-RNA concentration leads to one single RNA core, the intermediate concentration results in one central core surrounded by a single layer of smaller RNA speckles, and the lowest concentration yields DR-ACs with a fully speckled RNA phase. These results indicate that limited fusion of phase-separated RNA nuclei occurs prior to co-assembly with poly(A_20_-x)_n_ as the RNA concentration is lowered, resulting in kinetically trapped small RNA assemblies.^14^

**Figure 1.**
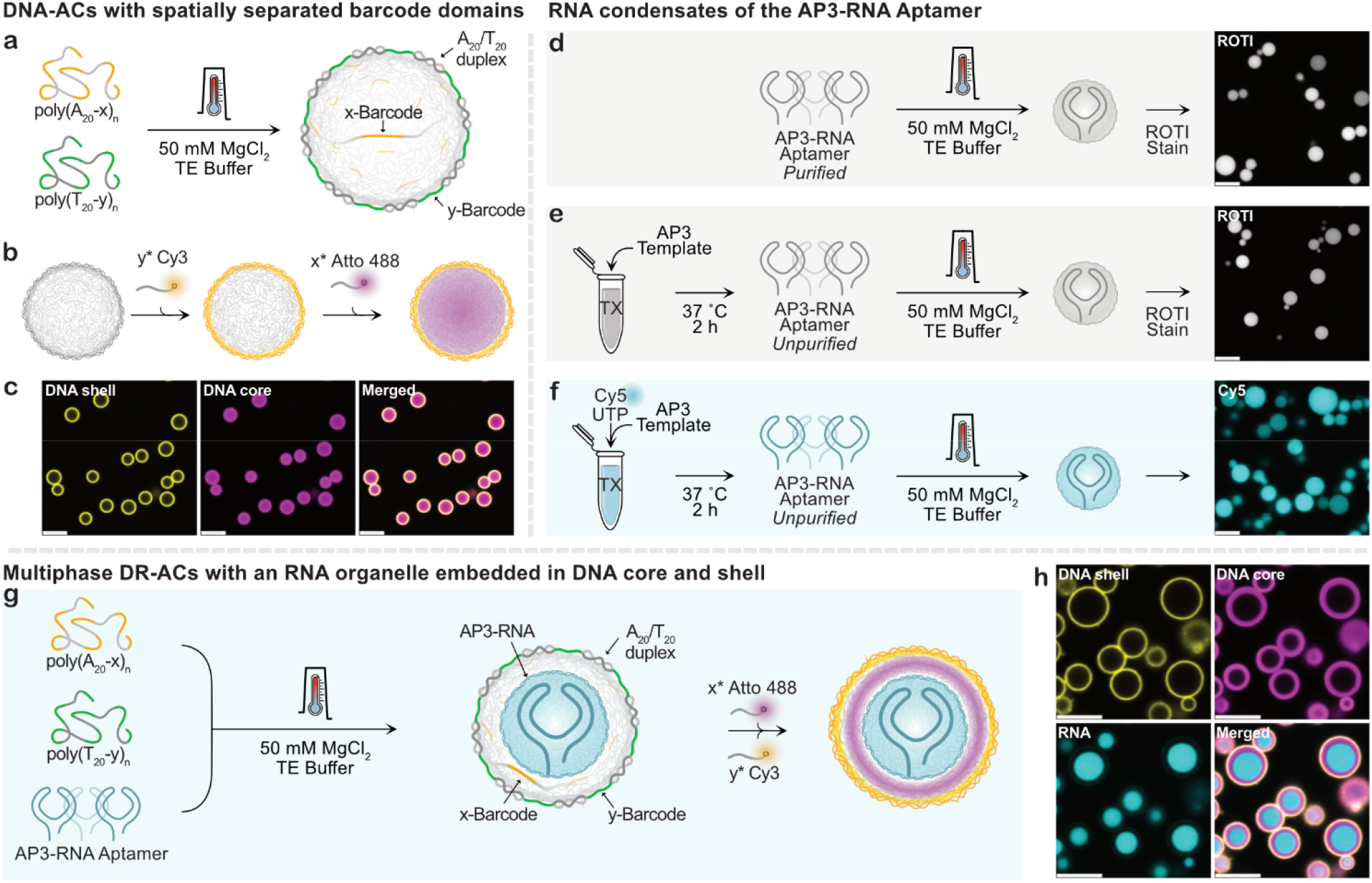
Synthesis of all-DNA ACs, RNA condensates and DNA–RNA artificial cells (DR-ACs) **a**) Scheme of DNA-AC formation from poly(A_20_-x)_n_ and poly(T_20_-y)_n_. **b**) The x and y barcodes enable post-assembly labeling of distinct DNA-AC regions. **c**) CLSM images of DNA-ACs (144 ng µL^-1^ poly(A_20_-x)_n_, 48 ng µL^-1^ poly(T_20_-y)_n_ (3:1 ratio), and 50 mM MgCl_2_) labeled with y*-Cy3 (DNA shell) and x*-Atto 488 (DNA core). **d-f**) Scheme and CLSM imaging of RNA condensates formed at 50 mM MgCl_2_ from (d) HPLC-purified AP3-RNA aptamer (719 ng µL^-1^), (**e**) unpurified AP3-RNA transcription mixture, and (**f**) unpurified fluorescently-labeled AP3-RNA transcription mixture (e-f, see Methods). CLSM (d-e) by staining with ROTI dye or (f) using co-transcribed Cy5-UTP. **g-h**) Scheme and corresponding CLSM of multiphase DR-AC formation from purified Cy5-labeled AP3-RNA, poly(A_20_-x)_n_, and poly(T_20_-y)_n_ (16:3:1 ratio) at 50 mM MgCl_2_. Labeling: x*-Atto 488 (DNA core), and y*-Cy3 (DNA shell). All CLSM images were obtained after a 30 min incubation in 40 mM HEPES, 125 mM KCl, and 5 mM MgCl_2_ at room temperature (RT). Scale bars: 5 µm.

**Figure 2.**
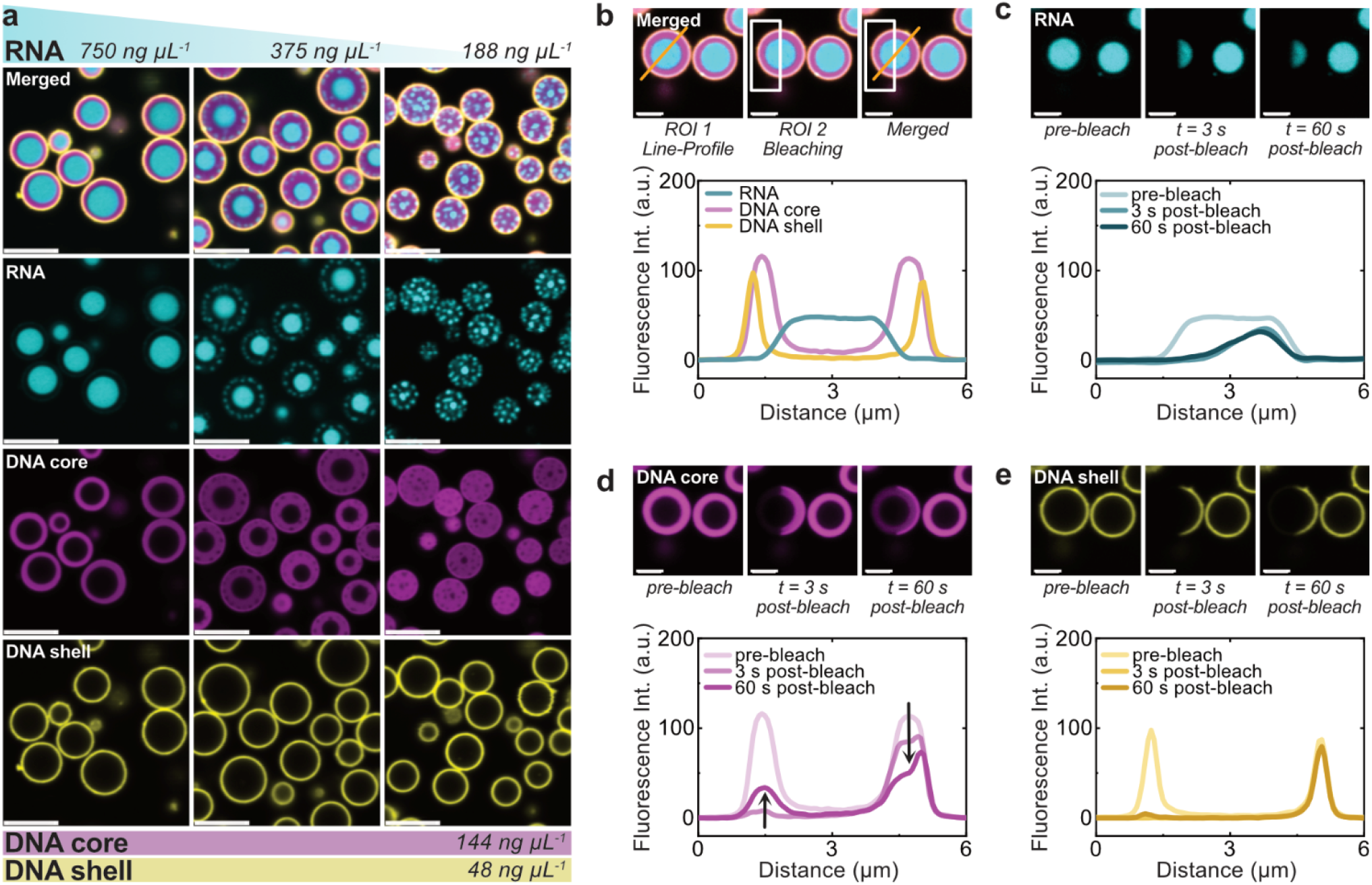
Structural and dynamic characterization of DR-ACs. **a**) CLSM images of DR-ACs prepared with varying concentrations of Cy5-labeled AP3-RNA (left: 750, middle: 375, right: 188 ng µL^-1^) and constant DNA concentrations (144 ng µL^-1^ poly(A_20_-x)_n_, 48 ng µL^-1^ poly(T_20_-y)_n_ (3:1 ratio), at 50 mM MgCl_2_. Labeling: x*-Atto 488 (DNA core) and y*-Cy3 (DNA shell). Scale bars: 5 µm. **b**) Representative DR-ACs (750 ng µL^-1^ Cy5-labeled AP3-RNA, 144 ng µL^-1^ poly(A_20_-x)_n_, 48 ng µL^- 1^ poly(T_20_-y)_n_ (16:3:1 ratio), and 50 mM MgCl_2_). Labeling: x*-Atto 488 (DNA core), and y*-Cy3 (DNA shell) prior to FRAP measurements. ROI1: cross-section of DR-AC for line profile. ROI2: bleaching region. Below: corresponding pre-bleach line profile. Pre-bleach, 3 s post-bleach, and 60 s post-bleach CLSM images of the RNA organelle (**c**), DNA core (**d**), and DNA shell (**e**), along with the corresponding line profiles. All CLSM images (panel a) were acquired, and FRAP experiments were performed after 2 h incubation in 40 mM HEPES, 125 mM KCl, and 5 mM MgCl_2_ at RT. Scale bars b-e: 2 µm.

To probe compartment dynamics, we performed fluorescence recovery after photobleaching (FRAP) analysis by hemispherical bleaching of a DR-AC and monitoring recovery along a line (**Figure 2b-e, Supplementary Video 1)**. The pre-bleach line cross-sectional profile reveals partial overlap between the phases, which stems from diffraction limitations in CLSM, but at the poly(A_20_-x)_n_–poly(T_20_-y)_n_ interface also from tight A_20_/T_20_ binding with an interdigitated interface. We recorded CLSM images immediately before bleaching and at 3 s and 60 s post-bleach. The RNA compartment exhibits no detectable fluorescence recovery over the observed time frame, confirming an arrested, glassy-like state, consistent with a percolated RNA network assisted by efficient Mg^2+^ bridging.^14^ In contrast, the poly(A_20_-x)_n_ DNA core within region of interest (ROI) 2 displays substantial fluorescence recovery and is accompanied by a corresponding decrease in fluorescence in the unbleached half, consistent with a more liquid-like character.^15^ Notably, the interfacial poly(A_20_-x)_n_–poly(T_20_-y)_n_ region (approximately at 5 µm in the corresponding line profile) shows reduced recovery in the poly(A_20_-x)_n_ channel (magenta) due to the aforementioned interdigitation and A_20_/T_20_ crosslinking at the interface. The shell composed of interfacially bonded poly(T_20_-y)_n_ correspondingly shows no recovery (yellow channel).

### Functionality of RNA organelles in DR-ACs

Next, we examine the extent to which functional RNA properties, such as those of aptamer binding, are maintained in the condensate state. The function of RNA aptamers critically depends on their complex secondary and tertiary structures, and subtle structural changes typically lead to loss of activity.^20^ We explore the retention of activity for the AP3-RNA aptamer that selectively binds to enhanced green fluorescent protein (eGFP)^23^ and for the so-called Spinach aptamer that binds the small molecule 3,5-difluoro-4-hydroxybenzylidene imidazolinone (DFHBI), subsequently inducing a conformational change that activates fluorescence.^25^ The AP3 and Spinach DR-ACs were created with identical assembly protocols from direct RNA transcription mixtures with 144 ng µL^-1^ poly(A_20_-x)_n_ and 48 ng µL^-1^ poly(T_20_-y)_n_ (3:1 ratio) at 50 mM MgCl_2_ (see Methods). Upon eGFP addition, the protein accumulates into AP3 DR-ACs within minutes, whereas the smaller DFHBI diffuses into Spinach DR-ACs within seconds and lights up upon binding (**Figure 3a,b**). To confirm aptamer-specific cargo recognition and exclude any non-specific binding, we also examined mixed populations of both ACs (AP3 DR-AC shell in yellow; Spinach DR-AC shell in magenta). Indeed, specific eGFP accumulation in AP3 DR-ACs occurs, while Spinach DR-ACs show no detectable protein recruitment (**Figure 3c; Supplementary Video 2**). Conversely upon addition of DFHBI, fluorescence activates only within Spinach DR-ACs and remains undetectable in AP3 DR-ACs (**Figure 3d**). While our observations cannot be extrapolated to all RNA aptamers, the successful functioning of both the protein-AP3 and the small-molecule-Spinach DR-ACs suggests that RNA aptamers may largely retain their selective binding functions when embedded in RNA condensate phases, even in those with arrested dynamics, and when confined in DNA shell condensates.

**Figure 3.**
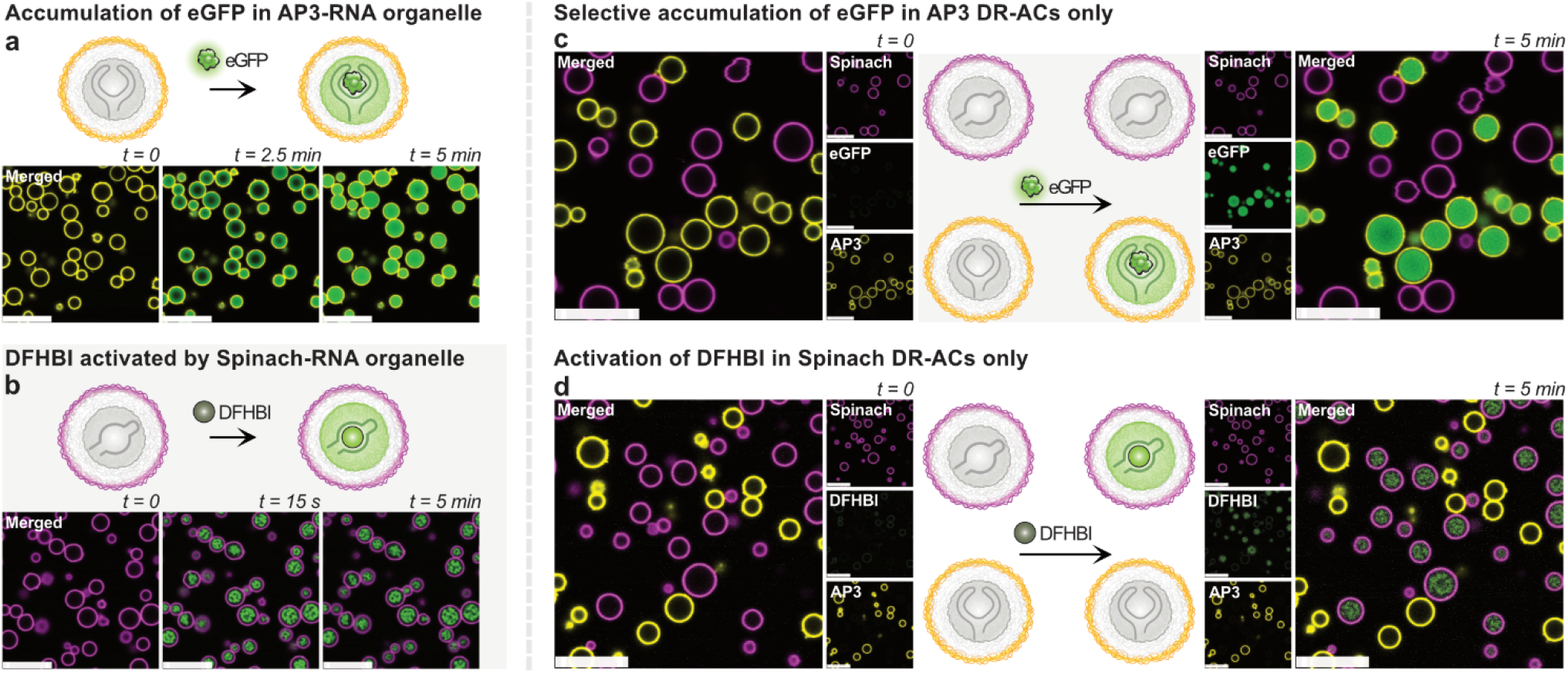
RNA aptamer functionality is maintained in condensate states. **a**) Scheme and CLSM of eGFP accumulation in AP3 DR-ACs. Conditions: AP3-RNA, poly(A_20_-x)_n_, and poly(T_20_-y)_n_ (3:1 ratio) DR-ACs and 1 µM eGFP; y*-Cy3 label (DNA shell); AP3-RNA used unpurified from transcription. **b**) Scheme and CLSM of DFHBI activation by Spinach DR-ACs. Conditions: Spinach-RNA, poly(A_20_-x)_n_, and poly(T_20_-y)_n_ (3:1 ratio) DR-ACs and 5 µM DFHBI; y*-DY-647P1 label (DNA shell); Spinach-RNA used unpurified from transcription. c-d) Competition experiments: Scheme and CLSM of selective (**c**) eGFP accumulation in AP3 DR-ACs (yellow shell) and (**d**) DFHBI activation by Spinach DR-ACs (magenta shell) by adding (c) 1 µM eGFP or (d) 5 µM DFHBI. All experiments were performed after 2 h incubation in 40 mM HEPES, 125 mM KCl, and 5 mM MgCl_2_ at RT. Scale bars: 10 µm.

### Serum-controlled protein release and enhancing serum stability

An important advantage of DNA-based ACs over RNA-based condensates is their enhanced stability in biologically relevant environments. RNA instability in serum arises primarily from susceptibility to RNase-mediated degradation, whereas DNA remains comparatively resistant. For instance, upon addition of 25% fetal bovine serum (FBS), the AP3-RNA organelles degrade within 90 s, allowing the fluid-like poly(A_20_-x)_n_ DNA core to occupy the vacated volume (**Figure 4a,b**). Time-lapse imaging and line-profile analysis confirm surface erosion of the RNA organelle (**Figure 4b–e, Supplementary Video 3**). This degradation can, however, be leveraged to achieve serum-triggered protein release via RNA degradation (**Figure 4f**). To this end, we exposed AP3 DR-ACs preloaded with eGFP to 25% FBS, resulting in rapid RNA degradation followed by gradual diffusion of eGFP out of the ACs (**Figure 4g,h; Supplementary Video 4**). These results demonstrate that DR-ACs can function not only as macromolecular capture platforms but also as serum-responsive release systems.

**Figure 4.**
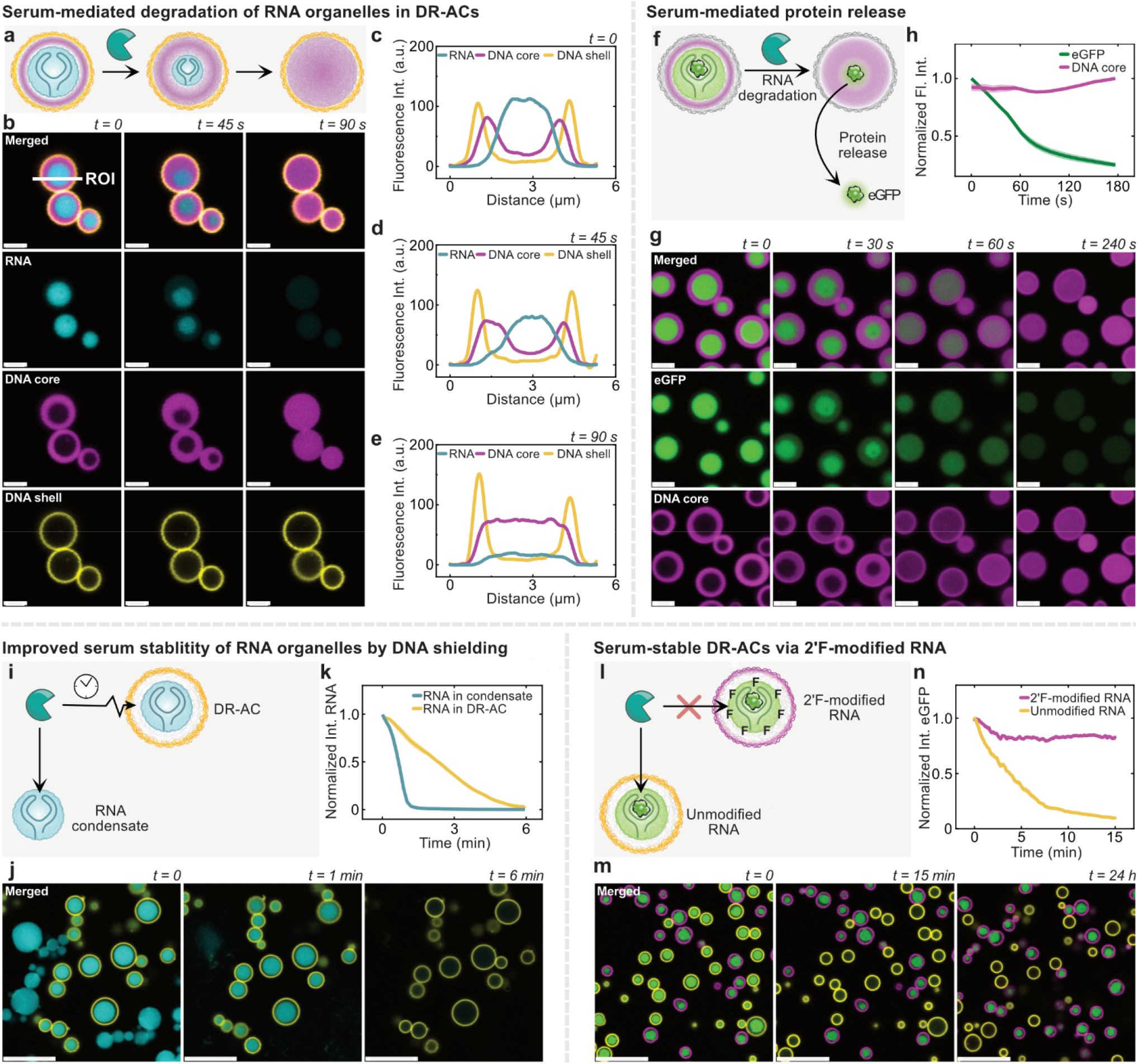
Serum-controlled protein release and serum stability. **a-b**) Scheme and CLSM of RNA organelle degradation within AP3 DR-ACs after 25% FBS addition. The RNA organelle degrades within 90 s, whereas the DNA remains intact. (DR-AC composition: Cy5-labeled AP3-RNA, poly(A_20_-x)_n_, and poly(T_20_-y)_n_ (16:3:1 ratio) DR-ACs; y*-Cy3 label (DNA shell); x*-Atto 488 label (DNA core)). **c-e**) Line profile analyses of ROI from panel b. **f-g**) Scheme and CLSM AP3 DR-AC-controlled eGFP release after 25% FBS addition. (DR-AC composition: AP3-RNA, poly(A_20_-x)_n_, and poly(T_20_-y)_n_ (11:4:1 ratio) DR-ACs; no visible poly(T_20_-y)_n_ shell labeling; x*-Atto 647N label (DNA core); pre-loaded with eGFP (final: 5 nM)). **h**) Average normalized fluorescence intensity of the poly(A_20_-x)_n_ DNA core and eGFP fluorescence. n = 5. **i-j**) Scheme and CLSM of the protective effect of DNA shield on RNA organelles upon 25% FBS addition for a 1:1 mixture of AP3-RNA condensates and AP3 DR-ACs. **k**) Corresponding fluorescence intensity plots of AP3 DR-ACs and AP3-RNA condensates. n = 8, 9. The RNA organelle in the DR-ACs (yellow) undergoes slower FBS-mediated degradation than unprotected RNA condensates (cyan). (Composition: AP3-RNA condensates: Cy3-labeled AP3-RNA. AP3 DR-AC: Cy3-labeled AP3-RNA, poly(A_20_-x)_n_, and poly(T_20_-y)_n_ (21:4:1 ratio); y*-DY-647P1 label (DNA shell)). **l-m**) Scheme and CLSM of the stabilization effect of 2’F-modifications in an RNA organelle in DR-ACs against FBS-mediated degradation for a 1:1 mixture of unmodified (yellow shell) and 2’F-modified (magenta shell) AP3 DR-ACs with captured eGFP after 25% FBS addition. **n**) Corresponding fluorescence intensity plots of eGFP fluorescence inside DR-ACs. n = 10. The eGFP is released from the unmodified RNA organelles, whereas 2’F-modified RNA organelles retain eGFP for 24 h. (Composition: Unmodified AP3 DR-AC: AP3-RNA, poly(A_20_-x)_n_, and poly(T_20_-y)_n_ (20:4:1 ratio). 2’F-modified DR-AC: 2’F-modified AP3-RNA, poly(A_20_-x)_n_, and poly(T_20_-y)_n_ (11:4:1 ratio). Labeling: unmodified DR-ACs: y*-Cy3 label (yellow, DNA shell); 2’F-modified DR-ACs: y*-DY-647P1 label (magenta, DNA shell), preloaded with 7 nM eGFP). Shaded areas in panels h), k) and n) (partly invisible as too small) represent the standard deviation. All experiments were performed after 1-2 h incubation in 40 mM HEPES, 125 mM KCl, and 5 mM MgCl_2_ at RT. Scale bars (b,g): 2 µm, (j,m): 10 µm.

To assess whether the DNA condensate shell has a protective effect on the RNA, we compare the stability of free AP3-RNA condensates with AP3-RNA organelles inside DR-ACs when exposed to 25% FBS (**Figure 4i-k; Supplementary Video 5**). While RNA condensates almost completely degrade within ∼75 s, the RNA organelles in DR-ACs degrade substantially slower (∼300 s). In addition to slower access to the RNA organelle, the degradation mechanism is also altered. Pure RNA condensates swell and dissolve, whereas RNA organelles in DR-ACs are rather surface eroded.

For applications requiring enhanced serum stability, we turned to 2’-fluoro (2’F) pyrimidine modifications inside the AP3-RNA aptamer to increase nuclease resistance.^26,27^ Such post-selection chemical changes to an RNA aptamer can perturb the folded structure, and thereby compromise target binding. Therefore, before incorporating the modified RNA into DR-ACs, we first assessed whether the 2’F AP3-RNA would retain eGFP binding. Remarkably, biolayer interferometry measurements reveal only a modest increase in the dissociation constant (*K*_D_) from 69 to 140 nM after modification (**Figure S6a-b**). This underscores that 2’F AP3-RNA largely retains eGFP recognition and that the electronegative 2’F pyrimidine substitution only minimally affects aptamer folding, binding, or both. Importantly, the modified aptamer also remains stable in solution (25% FBS) for at least 120 h (**Figure S6c-d**). Despite the altered chemical composition, 2’F-modified AP3-RNA forms condensates following the same temperature ramp protocol and RNA concentration (750 ng µL^-1^) as unmodified RNA and can be incorporated as organelle in DR-ACs by slightly reducing the final RNA and DNA concentrations (201 ng µL^-1^ 2’F-modified AP3-RNA, 72 ng µL^-1^ poly(A_20_-x)_n_, and 18 ng µL^-1^ poly(T_20_-y)_n_) and adjusting their ratios (11:4:1 ratio). This observation is particularly notable given that previous reports implicated the now partially replaced 2’-OH group in promoting RNA phase separation and stabilizing condensate formation.^28^

Consistent with enhanced nuclease resistance of the molecular 2’F-modified AP3-RNA, its RNA condensates and DR-ACs also exhibit minimal degradation in serum as shown in differential degradation experiments (**Figure 4l-n; Figure S7, S8, Supplementary Video 6, 7**). Most importantly, in mixed populations of unmodified and 2’F-modified AP3 DR-ACs preloaded with eGFP, only the unmodified RNA organelles degrade and release the cargo upon addition of 25% FBS, whereas 2’F-modified RNA organelles retain eGFP for at least 24 h. This demonstrates that chemical stabilization of RNA can be combined with phase separation procedures to furnish complex DR-AC structures and to achieve long-term serum stability.

### Sender–Receiver communication and integrated signal processing in single DR-ACs

Having established selective capture of protein cargo, we next investigate whether the same AC can actively program its environment to generate its own cargo. We first implement a multi-cell-like communication pathway between a DNA-AC acting as a Sender and a DR-AC acting as a Receiver, followed by integration of both functionalities within a single self-actuating DR-AC. The mechanism is built around DNA-triggered transcriptional gating inside the DNA part of the ACs, followed by RNA-triggered translational gating using a riboswitch in a cell-free transcription–translation (TX-TL) system to produce eGFP that is subsequently captured in the RNA organelle.^29,30^

To implement gated transcription within ACs, the x-barcode of the poly(A_20_-x)_n_ DNA core is exploited to position a DNA transcription template with a x*-P*-A*-x* sequence between two x barcodes. P* is the promoter binding site and A* is a flexible sequence acting as a template for the RNA transcript (A) of interest (**Figure 5a**).^29^ Functionalization proceeds by adding a stoichiometric amount of the template strand to the poly(A_20_-x)_n_ and poly(T_20_-y)_n_ mixture prior to AC formation. In the absence of promoter hybridization, RNA polymerase cannot initiate transcription, rendering the system transcriptionally dormant. To enable higher-level signal transduction, we introduce a second regulatory layer in the form of gated protein expression (GPE).^30^ As illustrated in **Figure 5b**, the mRNA encoding the protein output is translationally inactive through a defined secondary structure that occludes the ribosome binding site (RBS), thereby preventing ribosome association. An activator RNA strand (A), produced by transcription within the ACs from the A* domain, is required to bind to the folding region and restore RBS accessibility to initiate translation. Because the RBS itself is excluded from the folded region, the activator strand can be freely designed without sequence constraints.

**Figure 5.**
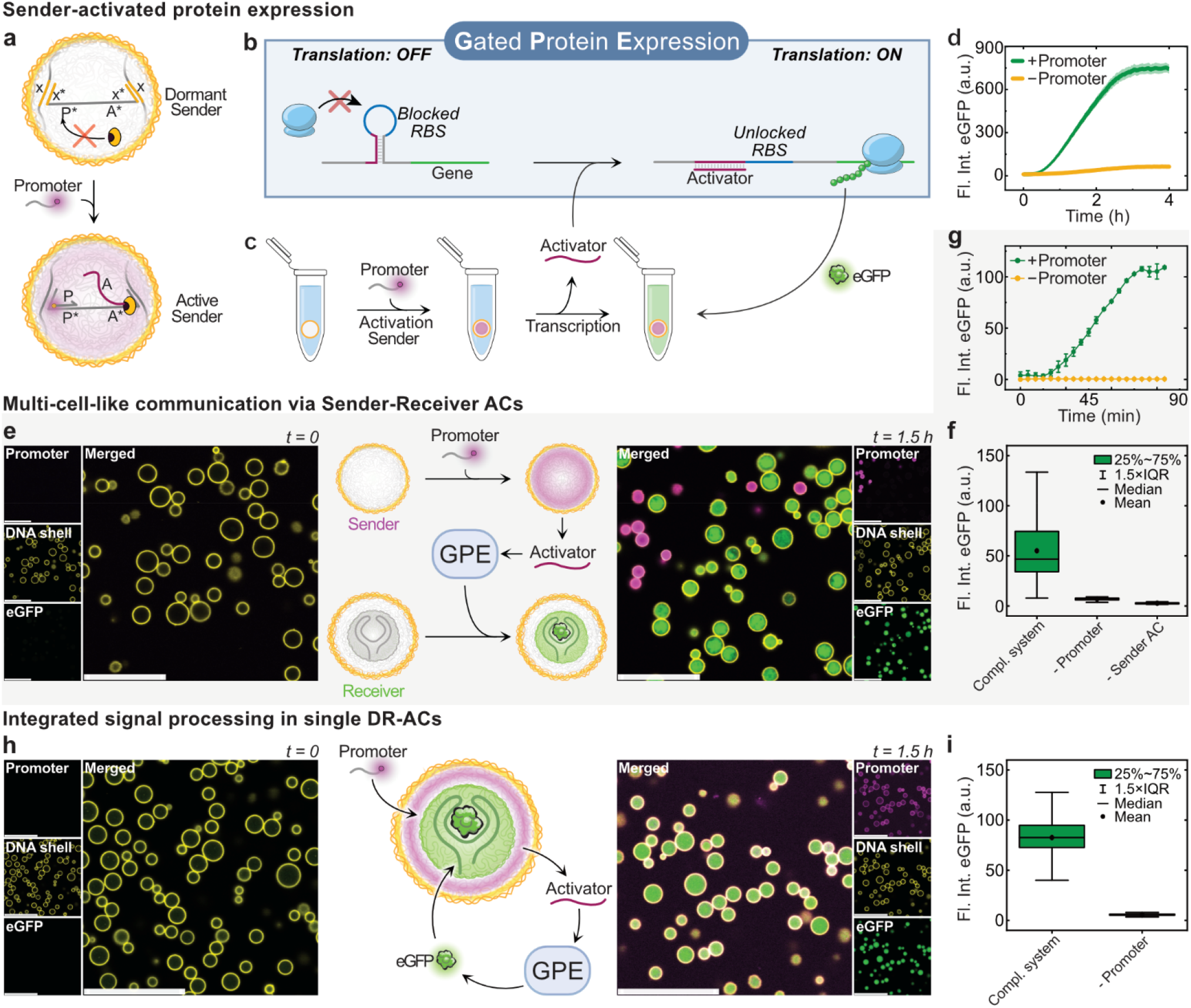
Sender–Receiver AC-AC communication and integrated signal processing in single DR-ACs. **a**) Scheme of a DNA-AC Sender containing a gated T7 transcription module. Transcription of the encoded RNA strand (A) occurs only upon hybridization of the promoter strand to the promoter binding site (P*). **b**) Scheme of gated protein expression (GPE), in which a defined mRNA secondary structure occludes the RBS, preventing translation. Hybridization of A disrupts the fold and enables the ribosome to initiate translation. **c**) Communication scheme: Promoter-mediated activation of the Sender AC induces transcription of A that diffuses out and triggers the GPE to yield eGFP. **d**) Real-time fluorescence measurements from the Sender AC-only system showing promoter-dependent eGFP production in solution, with minimal signal in the absence of the promoter (Conditions: Sender DNA-AC: poly(A_20_-x)_n_, poly(T20-y)_n_ (3:1 ratio), and 4.4 µM x*-P*-A*-x*) with or without 500 nM promoter, at 37 °C in TX-TL containing 12.5 ng µL^-1^ gated mRNA template). N = 3; shaded areas (partially invisible as too small) represent the standard deviation. **e**) CLSM images of the Sender–Receiver AC-AC system before and after promoter addition. Activated Sender ACs (magenta) transcribe A that diffuses and triggers the GPE to yield eGFP (green), which is selectively captured by the Receiver AC lighting up in green. (Conditions: Sender DNA-AC as in d; Receiver DR-AC (AP3-RNA, poly(A_20_-x)_n_, and poly(T_20_-y)_n_ (15:3:1 ratio) both ACs labeled with y*-Cy3; 37 °C, 1.5 h in TX-TL containing 12.5 ng µL^-1^ gated mRNA template, before and after addition of 500 nM DY-647P1-labeled promoter). **f**) Quantification of eGFP fluorescence per Receiver AC after 90 min with all components present (Compl. system), compared to negative controls lacking the promoter (-Promoter) or the Sender AC (-Sender). Data calculated from CLSM images with a minimum of 500 ACs per replicate (N = 3). Most Receiver ACs in the negative controls (‘-Promoter’ and ‘-Sender’) remain below the detection limit. **g**) Reaction kinetics of Sender–Receiver AC-AC communication, in the presence and absence of a promoter, where the average eGFP accumulation in Receiver ACs is monitored every 5 minutes with flow cytometry. Averages and standard deviations of N = 2 replicates are shown (reaction conditions equal to panel e, minus the y*-Cy3 label). **h**) Scheme and CLSM of integrated signal processing in single DR-ACs, in which promoter uptake (magenta) induces transcription of the Activator strand A that externally triggers the GPE, expressing eGFP that is subsequently captured within the RNA organelle of the same AC (Conditions: DR-AC: AP3-RNA, poly(A_20_-x)_n_, poly(T20-y)_n_ (15:3:1 ratio), and 1.0 µM x*-P*-A*-x*), labeled with y*-Cy3, with or without 500 nM promoter, at 37 °C in TX-TL containing 12.5 ng µL^-1^ gated mRNA template). **i**) Quantification of eGFP fluorescence per DR-AC after 90 min with all components present (Compl. system), or in the absence of the promoter (-Promoter), demonstrating robust promoter-dependent self-actuation. Data calculated from CLSM images with a minimum of 400 ACs per replicate (N = 3). Scale bars: 20 µm.

We first focus on validating GPE from gated transcription from Sender DNA-ACs when immersed in the surrounding TX-TL environment. Upon promoter addition, transcription of the activator A is initiated inside the Sender DNA-ACs (**Figure 5c**). Plate reader fluorescence measurements show promoter-dependent eGFP production within ∼30 min, reaching a maximum after approximately 3 h (**Figure 5d**). In the absence of the promoter, only negligible fluorescence is observed, confirming high dormancy of both transcriptional and translational gates when combined.

This system is next extended to establish multi-cell-like communication between Sender and Receiver ACs. To this end, we mixed Sender DNA-ACs with an equal volume of AP3 DR-ACs acting as Receiver ACs, all immersed in a TX-TL medium. **Figure 5e** shows that addition of a fluorescently labeled promoter selectively identifies the Sender ACs, which turn magenta, where transcription of the activator RNA (A) is triggered. This activator RNA strand then diffuses into the TX-TL environment, activates the GPE, and the produced eGFP selectively accumulates within the RNA organelles of the Receiver ACs due to capture by the AP3 aptamer. At *t* = 0, the eGFP fluorescence of the Receiver AC remains undetectable (RFU < 0.1) in CLSM images. Quantification after 90 min reveals a mean Receiver AC fluorescence of 55.0 RFU when all components are present (Compl. system) compared to 2.7 RFU in the absence of a Sender AC (-Sender). The second negative control combines gated translation and transcription modules in the absence of the promoter (-Promoter) and has a slightly elevated signal of 6.8 RFU (**Figure 5f**). Flow cytometry detects eGFP in Receiver ACs after ∼15 min and plateaus after ∼90 min (**Figure 5g**). The plateau in the flow cytometry signal as compared to the longer increase in the plate reader experiment lacking the Receiver AC (**Figure 5d**) stems from efficient accumulation of eGFP into the Receiver ACs and saturation of binding sites. Any additional eGFP produced in solution therefore remains undetected by flow cytometry.

Finally, we integrated Sender and Receiver functionalities into a single self-actuating DR-AC. In this configuration, promoter uptake into the poly(A_20_-x/x*P*A*x*) DNA core of the DR-AC initiates activator (A) transcription, which similarly triggers the GPE in the external TX-TL environment, and finally the translated eGFP accumulates within the AP3-RNA organelles in the same AC (**Figure 5h**). CLSM quantification after 90 min (**Figure 5i**) shows robust promoter-dependent self-actuation, with a mean fluorescence intensity of 83.6 RFU compared to 5.6 RFU without promoter (negative control).

Collectively, these results demonstrate that programmable DNA host ACs combined with functional RNA aptamer organelles enable multi-cell-like communication as well as integrated signal processing within single DR-ACs. The absence of sequence constraints at both transcriptional and translational levels enables potential for extension toward multiplexed protein outputs. When combined with universally triggered transcription strategies such as our recently discovered T7-Lock system^29^, this platform provides a powerful framework for higher-level biochemical computation with spatial compartmentalization and functional outputs.

## Conclusion

In this work, we have established multiphase DNA–RNA artificial cells (DR-ACs) that integrate an RNA condensate organelle within a programmable DNA core–shell AC. Using a one-pot thermal assembly strategy, we generate tunable three-phase architectures, in which a glassy-like RNA organelle is embedded within a liquid-like DNA core and enclosed by a crosslinked, but permeable DNA shell. The DNA/RNA ratios effectively regulate RNA organelle size and size distribution, while the shell simultaneously provides a temporal protective element against serum-mediated RNA degradation.

Beyond structural organization, DR-ACs translate compartmentalization into function. The embedded RNA organelles preserve their aptamer-encoded functions, enabling protein accumulation or small-molecule activation – a phenomenon that is very beneficial and can potentially be exploited for the controlled delivery and release of drugs or biologics. In biologically relevant media, the nuclease sensitivity of RNA becomes an operational feature: RNases in serum digest the pre-loaded RNA organelles, effectively releasing their proteins, while the DNA-AC scaffold remains largely intact. Conversely, incorporation of 2’-fluoro modifications in the RNA extends RNA organelle lifetimes in serum without substantially compromising aptamer activity or DR-AC formation.

Finally, we coupled transcriptional signaling localized within the DNA core to external cell-free reaction networks, such that RNA signals generated inside ACs trigger translation in the surrounding environment and the resulting protein products are selectively concentrated within the RNA organelles. This design enables Sender–Receiver communication across AC populations as well as self-actuating signal processing within single DR-ACs.

Collectively, this work provides a framework for unifying structural programmability and biochemical function within multiphase DNA-RNA hybrid condensates, advancing synthetic systems a step further toward the spatial organization characteristics of living cells. DR-ACs combine structurally addressable DNA compartments with functional RNA organelles that enables controlled partitioning, localized biochemical activity, and inter-compartment communication. As such, they establish a platform for higher-order synthetic biology systems that more closely emulate cellular organization and signaling.

## Author Information

### Supporting Information

The supporting information is available free of charge at <u>XXXX</u>.

– Materials and instrumentation, experimental details, data analysis methods, and Figures (pdf)
– Supplementary Video 1: FRAP on multiphase DNA–RNA artificial cells
– Supplementary Video 2: Selective accumulation of eGFP in AP3 DR-ACs
– Supplementary Video 3: Serum-triggered RNA organelle degradation in DR-ACs
– Supplementary Video 4: Serum-triggered eGFP release from AP3 DR-ACs
– Supplementary Video 5: Improved serum stability of RNA organelle by DNA shielding
– Supplementary Video 6: Serum-stable 2’F-modified RNA condensates
– Supplementary Video 7: Serum-stable 2’F-modified RNA organelles in DR-ACs

## Acknowledgment

We thank Dr. Christoph Drees, Dr. Brigitta Duzs, Rune Kersebaum, and Sebastian Bauer for discussions. We acknowledge support by the EU in the framework of the ERC Consolidator Grant to AW – M3ALI (101001638). D.H. and A.W. acknowledge support from the German Research Foundation (Grant WA 3084/19-1). AW acknowledges funding via a Gutenberg Research Professorship underpinning his Life-Like Materials Program. J.V. and L.C. thank the support of Novo Nordisk Foundation (Grant No. NNF22OC0080492), and the VILLUM Foundation (Grant No. Villum Investigator Grant 6960). The authors gratefully acknowledge the IMB Flow Cytometry Core Facility for their support and assistance in this work.

