## Supplementary Information for "RNA Organelles in DNA-based Artificial Cells Provide Spatial Aptamer Functions and Enhanced Signal Processing"

### 1. Materials

HEPES (≥99.5%), n-hexadecane (99.0%), potassium chloride (99%), magnesium chloride solution (BioUltra, 2 M in $H_2O$), Tween®20, BSA heat shock fraction (≥98%), FBS superior, and 96-well black flat-bottom plates were purchased from Sigma Aldrich. ROTIPHORESE®50x TAE buffer and ROTI®GelStain (20,000x conc) were purchased from Carl Roth. GeneRuler 50 bp DNA ladder, Ultra Low Range dsDNA Ladder, 1× TE Buffer (10 mM Tris-HCl (pH 8.0), 0.1 mM EDTA), UltraPure™ Glycerol, UltraPure™ Agarose, dithiothreitol (100 mM), Proteintech ChromoTek eGFP, recombinant purified protein (1 µg $µL^{-1}$), and Phusion™ High-Fidelity DNA Polymerase (2U/µL) were obtained from Thermo Fisher Scientific. 5Z-[(3,5-difluoro-4-hydroxyphenyl)methylene]-3,5-dihydro-2,3-dimethyl-4H-imidazol-4-one (98%, DFHBI) from Hycultec, as well as aminoallyl-UTP-Cy3 (1 mM), aminoallyl-UTP-ATTO-647N (1 mM), aminoallyl-UTP-Cy5 (1 mM), and ATP and GTP (100 mM) from Jena Bioscience. Cy5-UTP (10 mM) was purchased from APExBIO. 2′F-dCTP and 2′F-dUTP (100 mM) were purchased from Metkinen Chemistry. Agarose, hydrochloric acid (0.1 N, Titripur), and sodium chloride (≥98%, technical) were purchased from VWR. T7 RNA Y639F polymerase (100x) was obtained from Aptamist ApS. All chemicals were used as received.

GeneJET PCR purification kit was purchased from Thermo Fisher Scientific and RNA Clean & Concentrator-25 kit was obtained from Zymo Research. Octet Ni-NTA Biosensors were purchased from Sartorius.

HiScribe® T7 Quick High Yield RNA Synthesis Kit (T7 Polymerase Mix and NTP Buffer Mix, E2050L), PURExpress® *In Vitro* Protein Synthesis Kit (E6800L), Monarch® Spin RNA Isolation Kit (T2110S). *Taq* polymerase (5,000 units/mL),10× Standard *Taq* Buffer (100 mM Tris-HCl, 500 mM KCl, 15 mM $MgCl_2$, pH 8.3 @25 °C), RNase Inhibitor Murine (40,000 units/mL (M0314S), T4 DNA ligase (400,000 units/mL), 10× T4 DNA ligase buffer (500 mM Tris-HCl, 100 mM $MgCl_2$, 10 mM ATP, 100 mM DTT, pH 7.5 @25 °C), Exonuclease I (40 U $µL^{-1}$), Exonuclease III (200 U $µL^{-1}$), phi29 (Φ29) DNA polymerase (10 U $µL^{-1}$), 10× phi29 Reaction Buffer (500 mM Tris-HCl, 100 mM $MgCl_2$, 100 mM $(NH_4)_2SO_4$, 40 mM DTT (pH 7.5 @25 °C) inorganic Pyrophosphatase (E.coli) (0.1 U $µL^{-1}$), deoxynucleotides (dNTP) solution mix (10 mM each), and nuclease-free water were purchased from New England Biolabs, and used according to the manufacturer's protocol (unless stated differently).

A DNA sequence containing the sequence for gated eGFP mRNA (T7 promoter, gated loop, RBS and eGFP gene, Table S1) was synthesized and subcloned into the pUC57 backbone by GenScript Biotech (Netherlands) B.V. The plasmid was delivered as a sequence-verified construct and used without further modification. Plasmid pIRES2-EGFP-p53 tethered (catalog #49245) was purchased from Addgene. AP3 DNA template, AP3 primer (forward) and AP3 primer (reverse) were obtained from Integrated DNA Technologies (IDT). All other oligonucleotides (with sequences listed in Table S2) were purchased from Biomers GmbH and dissolved in nuclease-free water. All buffers and DNA/RNA stock solutions, were dissolved in nuclease-free water as well. The 10× HEPES salt buffer (400 mM HEPES, 1.25 M KCl and 50 mM $MgCl_2$, pH 7.4 @ 25 °C) was prepared in-house.

### 2. General Characterization Methods & Instruments

**Instruments**

Oligonucleotide concentrations were determined with a DS-11 Spectrophotometer (DeNovix). Incubations and annealing were carried out on an Eppendorf ThermoMixer C with heated lid. Centrifugation was performed on a Sigma 1-16K centrifuge (Sigma Laborzentrifugen GmbH). Confocal laser scanning microscopy (CLSM) was performed on a Leica Stellaris 5 microscope with a 63× oil-immersion objective.

All heating ramps were performed on a qTOWER$^3$ Real-time Quantitative PCR (Analytik Jena). Standard heating ramp: samples were heated from 20 °C to 95 °C (8.0 °C $s^{-1}$), held at 95 °C for 180 s (unless stated differently), and rapidly cooled to 4 °C (6.0 °C $s^{-1}$), where they were held for ≥5 min (heated lid: 100 °C).

Real-time fluorescence measurements were recorded on a TECAN (SPARK control v3.1) microplate reader with excitation and emission wavelengths at 485 and 510 nm and a bandwidth of 10 nm. All samples were prepared in a volume of 20 µL, placed in a Corning® 384-well clear-bottom polystyrene plates with a non-binding surface, and covered with 10 µL hexadecane and measured every 60 s for 4 h at 37 °C.

Flow cytometry was performed on a Novocyte Quantean, Agilent (Acquisition software: NovoExpress v.1.6.2) equipped with 4 excitation lasers (violet 405 nm, blue 488 nm, yellow-green 561 nm and red 640 nm), 16 fluorescence detectors, and an autosampler. Data analysis was performed using FlowJo (v.11.1.0).

Bio-layer interferometry (BLI) kinetic studies were conducted using an OctetRed96 instrument from ForteBio. All binding experiments were performed at an orbital shaking speed of 1,000 rpm. Data were analyzed using GraphPad Prism (v.5.0). Agarose gels were imaged in a Gel Doc™ EZ Imager from BIO-RAD.

**Image analysis**

*Stability of ACs in serum:* Time-lapse CLSM images were processed and analyzed using Leica LAS X software (v4.3.0.24308). Identical gain and laser power were used within each experimental dataset. The regions of interest (ROIs) were manually defined around/within individual particles of interest, of which the number is specified in the corresponding figure captions. Within each ROI, the mean fluorescence intensity was determined for each frame within the time-lapse series and plotted as a function of time to evaluate particle stability under serum-containing conditions.

*eGFP uptake in Sender-Receiver and integrated signal processing in single DR-ACs*: CLSM images at $t = 0$ and $t = 90$ min were processed and analyzed using Leica LAS X software (v4.3.0.24308). After threshold-based segmentation, objects were filtered (minimum size: 1000 pixels, roundness ≥ 0.95, exclusion of particles touching image borders). The mean fluorescence intensity of each resulting segmented object was extracted, and the average intensities and standard deviations per dataset were calculated.

*Particle diameter and count*: CLSM images were analyzed using Fiji (ImageJ, x64), where they were first subjected to a median filter (2-pixel radius) to reduce noise while maintaining particle boundaries. The automatic thresholding function was applied to generate binary masks. Closely adjacent objects were separated using a watershed algorithm. Quantitative and qualitative analysis was performed using the 'Analyze Particles' function (size range: 0.01 to infinity, circularity: 0.6-1.0, exclusion of particles touching the image borders). The Feret diameter and particle count per image were extracted.

**Data processing**

The plotted curves represent a mean of the measurements listed in the figure captions and methods section below. The error bars or shaded regions around the curve represent standard deviations calculated in OriginPro 2023.

### 3. Methods

**Synthesis of poly($A_{20}$-x)$_n$ and poly($T_{20}$-y)$_n$ ssDNA.** Long ssDNA strands were synthesized by first producing circular DNA templates, which were thereafter copied into long ssDNA chains with multiple repeating units by rolling circle amplification (RCA). This protocol is based on our previous report.[1]

*Synthesis of circular DNA templates.* In a total volume of 100 µL, 1 µM linear template (*Template ($A_{20}$-x) or Template ($T_{20}$-y)),* 1 µM of its corresponding ligation strand (*x ligation or y ligation, respectively*) and 100 mM NaCl were assembled in TE buffer. The resulting solutions were heated for 5 min at 85 °C and subsequently cooled to 20 °C at a rate of 1 °C min$^{-1}$. After annealing, to both tubes, 20 µL 10× T4 Ligase Buffer, 10 µL T4 Ligase (4000 U), and 70 µL nuclease-free water were added, resulting in a 200 µL reaction volume. After gentle mixing (10 min at 400 rpm), the samples were left to react at room temperature (RT) for 4 h, followed by a 20 min denaturation step at 70 °C. After cooling, 15 µL Exonuclease I (40 U µL$^{-1}$) and 15 µL Exonuclease III (200 U µL$^{-1}$) were added to each sample, after which they were incubated at 37 °C for 16 h under gentle mixing (300 rpm). Afterward, the samples were heated for 40 min at 80 °C to deactivate the exonucleases. The remaining circular templates were purified by filtration with Amicon Ultracentrifugal filters (10 kDa MWCO cutoff, Merck Millipore), followed by a 400 µL TE buffer washing step, which was repeated 3 times. The concentrations of the templates were determined on the spectrophotometer and adjusted to 2.5 µM with TE buffer. The templates were stored at -20 °C until further use.

*RCA synthesis of poly($A_{20}$-x)$_n$ and poly($T_{20}$-y)$_n$ ssDNA polymers.* In 200 µL PCR tubes, 25 µL dNTP solution mix (10 mM each), 20 µL pyrophosphatase (0.1 U µL$^{-1}$), 4 µL phi29 (Φ29) DNA polymerase (10 U µL$^{-1}$), 20 µL 10× phi29 Reaction Buffer, 4 µL 2.5 µM circular ($A_{20}$-x) or ($T_{20}$-y) template, along with 2 µL of their complementary primer (10 µM) (primer-x or primer-y, respectively) were added. The reaction volume was adjusted to 200 µL by the addition of 125 µL nuclease-free water. The reaction mixture was incubated for 60 h at 30 °C, followed by a 15 min cleaving step at 95 °C. The remaining ssDNA polymers were purified by filtration with Amicon Ultracentrifugal filters (30 kDa MWCO cutoff, Merck Millipore), followed by a 400 µL TE buffer washing step, which was repeated 3 times. The obtained poly($A_{20}$-x)$_n$ and poly($T_{20}$-y)$_n$ chains had concentrations of 3600 and 4200 ng µL$^{-1}$, respectively, and n ranging roughly from 10 to 100, averaging at ~37 repeating units.[2] The samples were diluted to 500 ng µL$^{-1}$ with TE buffer, and stored at -20 °C until further use.

*DNA-AC pre-mixes.* Five different types of poly($A_{20}$-x)$_n$ and poly($T_{20}$-y)$_n$ ssDNA pre-mixes were made in TE buffer. A): 231 ng µL$^{-1}$ poly($A_{20}$-x)$_n$ and 77 ng µL$^{-1}$ poly($T_{20}$-y)$_n$. B): 231 ng µL$^{-1}$ poly($A_{20}$-x)$_n$ and 0 ng µL$^{-1}$ poly($T_{20}$-y)$_n$. C): 0 ng µL$^{-1}$ poly($A_{20}$-x)$_n$ and 77 ng µL$^{-1}$ poly($T_{20}$-y)$_n$. D): 0 ng µL$^{-1}$ poly($A_{20}$-x)$_n$ and 100 ng µL$^{-1}$ poly($T_{20}$-y)$_n$. E): 231 ng µL$^{-1}$ poly($A_{20}$-x)$_n$ and 57 ng µL$^{-1}$ poly($T_{20}$-y)$_n$. All pre-mixes were thermally cleaved at 95 °C for 15 min in the ThermoMixer, after which they were stored at -20 °C until further use.

**Preparation of pure DNA-ACs *(Figure 1c)*.** A 200 µL PCR tube containing 2.5 µL DNA-AC pre-mix A, 0.5 µL 400 mM $MgCl_2$, and 1 µL TE buffer was subjected to the standard heating ramp (see above). Afterward, 2 µL y*-Cy3 label (6.25 µM) and 2 µL x*-Atto 488 label (6.25 µM) were added and left to hybridize at RT for at least 30 min. The resulting AC was gently vortexed, after which 2 µL was added to a 384-well glass-bottom plate (Corning) containing 28 µL 1× HEPES salt buffer. The condensates were left to sediment for at least 30 min before CLSM imaging.

**Preparation of pure RNA condensates *(Figure 1d-f)*.**

*Purified RNA*: A 200 µL PCR tube containing 2.5 µL TE buffer, 0.5 µL 400 mM $MgCl_2$, and 1 µL 100 µM AP3-RNA (purified, 2875 ng µL$^{-1}$, Biomers) was subjected to the standard heating ramp. Afterward, the resulting dispersion was gently vortexed, and 2 µL was added to a 384-well glass-bottom plate (Corning) containing 28 µL 1× HEPES salt buffer and 10× ROTI®GelStain. The condensates were left to sediment for at least 30 min before CLSM imaging.

*Unpurified RNA*: A 200 µL PCR tube containing 2.5 µL TE buffer, 0.5 µL 400 mM $MgCl_2$, and 1 µL unpurified transcript (Transcription: 1 µL AP3 template (25 µM), 1 µL T7 promoter (25 µM), 6.67 µL NTP Buffer mix, 1.33 µL T7 RNA Polymerase mix, and 10 µL nuclease-free water were gently homogenized and left to transcribe for 2 h at 37 °C), was subjected to the standard heating ramp. Afterward, the resulting dispersion was gently vortexed, and 2 µL was added to a 384-well glass-bottom plate (Corning) containing 28 µL 1×

HEPES salt buffer and 10× ROTI®GelStain. The condensates were left to sediment for at least 30 min before CLSM imaging.

*Unpurified Cy5-labeled RNA*: A 200 µL PCR tube containing 2.5 µL TE buffer, 0.5 µL 400 mM $MgCl_2$, and 1 µL unpurified transcript (Transcription: 1 µL AP3 template (25 µM), 1 µL T7 promoter (25 µM), 2 µL 0.1 mM aminoallyl-UTP-Cy5, 6.67 µL NTP Buffer mix, 1.33 µL T7 RNA Polymerase mix, and 8 µL nuclease-free water were gently homogenized and left to transcribe for 2 h at 37 °C) was subjected to the standard heating ramp. Afterward, the resulting dispersion was gently vortexed, after which 2 µL was added to a 384-well glass-bottom plate (Corning) containing 28 µL 1× HEPES salt buffer. The condensates were left to sediment for at least 30 min before CLSM imaging.

**Preparation of DR-ACs *(Figure 1g,h)*.** A 200 µL PCR tube containing 2.5 µL DNA-AC pre-mix A, 0.5 µL 400 mM $MgCl_2$, and 1 µL purified Cy5-labeled AP3-RNA (3000 ng $\mu L^{-1}$) (Transcription: 2 µL AP3 template (25 µM), 2 µL T7 promoter (25 µM), 4 µL 0.1 mM Aminoallyl-UTP-Cy5, 13.3 µL NTP Buffer mix, 2.67 µL T7 RNA Polymerase mix, and 16 µL nuclease-free water were gently homogenized and left to transcribe for 2 h at 37 °C, after which it was purified with a Monarch Spin RNA Isolation Kit and adjusted to 3000 ng $\mu L^{-1}$ with nuclease-free water), was subjected to the standard heating ramp. Afterward, 2 µL y*-Cy3 label (6.25 µM) and 2 µL x*-Atto 488 label (6.25 µM) were added and left to hybridize at RT for at least 30 min. The resulting dispersion was gently vortexed, after which 2 µL was added to a 384-well glass-bottom plate (Corning) containing 28 µL 1× HEPES salt buffer. The condensates were left to sediment for at least 30 min before CLSM imaging.

**Preparation of DR-ACs from RNA and either poly($A_{20}$-x)$_n$ or poly($T_{20}$-y)$_n$ *(Figure S2)*.** A 200 µL PCR tube containing 2.5 µL DNA-AC pre-mix B (poly($A_{20}$-x)$_n$ only) or C (poly($T_{20}$-y)$_n$ only), 0.5 µL 400 mM $MgCl_2$, 0.5 µL TE buffer, and 0.5 µL unpurified transcript (Transcription: 1 µL AP3 template (25 µM), 1 µL T7 promoter (25 µM), 2 µL 0.1 mM Aminoallyl-UTP-Cy5, 6.67 µL NTP Buffer mix, 1.33 µL T7 RNA Polymerase mix, and 8 µL nuclease-free water were gently homogenized and left to transcribe for 2 h at 37 °C), was subjected to the standard heating ramp. Afterward, 2 µL y*-Cy3 label (6.25 µM) (poly($T_{20}$-y)$_n$ only) or 2 µL x*-Atto 488 label (6.25 µM) (poly($A_{20}$-x)$_n$ only) were added and left to hybridize at RT for at least 30 min. The resulting dispersion was gently vortexed, after which 2 µL was added to a 384-well glass-bottom plate (Corning) containing 28 µL 1× HEPES salt buffer. The condensates were left to sediment for at least 30 min before CLSM imaging.

**Effect of poly($T_{20}$-y)$_n$ DNA shell concentration on RNA condensate size distribution and homogeneity *(Figure S3)*.** From DNA-AC pre-mix D (100 ng $\mu L^{-1}$ poly($T_{20}$-y)$_n$), stock solutions of 100, 50, 25, 12.5, 6.25, 3.13, and 0 ng $\mu L^{-1}$ poly($T_{20}$-y)$_n$ were made in nuclease-free water. In separate 200 µL PCR tubes, 2 µL of each poly($T_{20}$-y)$_n$ stock was added to 1 µL purified Cy5-labeled AP3-RNA (3000 ng $\mu L^{-1}$; synthesis protocol above), 0.5 µL 400 mM $MgCl_2$, and 0.5 µL TE buffer. The samples were subjected to the standard heating ramp. Afterward, 2 µL y*-Atto 488 label (6.25 µM) was added to each sample and left to hybridize at RT for at least 30 min. The resulting dispersions were gently vortexed, after which 2 µL was added to a 384-well glass-bottom plate (Corning) containing 28 µL 1× HEPES salt buffer. The condensates were left to sediment for at least 30 min before CLSM imaging. The particle diameter and count were analyzed using Fiji.

**Influence of plateau time in temperature ramp on RNA condensate size and size distribution *(Figure S4)*.** In separate 200 µL PCR tubes, 11 identical samples were made with 1 µL DNA-AC pre-mix D (100 ng $\mu L^{-1}$ poly($T_{20}$-y)$_n$), 1 µL purified Cy5-labeled AP3-RNA (3000 ng $\mu L^{-1}$; synthesis protocol above), 0.5 µL 400 mM $MgCl_2$, and 1.5 µL TE buffer. The samples were heated from 20 °C to 95 °C (8.0 °C $s^{-1}$), held at 95 °C for varying plateau times of (Δs) 15, 30, 60, 90, 120, 150, 180, 210, 240, 270 and 300 seconds, and cooled down to 4 °C (6.0 °C $s^{-1}$). Afterward, 2 µL y*-Atto 488 label (6.25 µM) was added to each sample and left to hybridize at RT for at least 30 min. The resulting dispersions were gently vortexed, after which 2 µL was added to a 384-well glass-bottom plate (Corning) containing 28 µL 1× HEPES salt buffer. The condensates were left to sediment for at least 30 min before CLSM imaging. The particle diameter and count were analyzed using Fiji.

**Synthesis of long transcription template [H].** A T7 promoter sequence was introduced to a sequence extracted from plasmid pIRES2-EGFP-p53 tethered by means of PCR. The PCR sample was prepared in 20-fold and consisted of 2 ng pIRES2-EGFP-p53 tethered plasmid, 0.2 µM Reverse Primer DNA, and 0.2

µM Forward Primer DNA (listed in Table S2). To that, 0.2 mM dNTPs, 1× *Taq* buffer (10 mM Tris-HCl, 50 mM KCl, 1.5 mM $MgCl_2$) and 2.5 units *Taq* polymerase were added. The sample volumes were adjusted to 100 µL each by addition of nuclease-free water. The PCR was performed on a qTOWER$^3$ Real-time Quantitative PCR (Analytik Jena) as follows: initial denaturation at 95 °C for 1 min, followed by 30 cycles of denaturation at 95 °C for 30 seconds, annealing at 58 °C for 40 seconds and extension at 68 °C for 90 seconds. This was followed by one final extension at 68 °C for 5 min after which all replicates were pooled. Template purification was performed by filtration with Amicon Ultracentrifugal filters (100 kDa MWCO cutoff, Merck Millipore), followed by a 500 µL TE buffer wash, and a double 500 µL nuclease-free water washing step. The concentration was determined with the spectrophotometer and diluted to 125 ng µL$^{-1}$ with nuclease-free water.

**RNA condensates and DR-ACs with varying RNA sequences *(Figure S5).*** Several types of RNA with varying sequences were transcribed from different DNA templates (DNA templates A-H (synthesis template H described above) are indicated in Table S2). Transcription: 2 µL DNA template A-G (25 µM) or DNA Template H (125 ng µL$^{-1}$), 2 µL T7 promoter (25 µM), 6 µL 0.1 mM Aminoallyl-UTP-Cy3, 13.3 µL NTP Buffer mix, 2.67 µL T7 RNA Polymerase mix, and 14 µL nuclease-free water were gently homogenized and left to transcribe for 2 h at 37 °C. Afterward, the transcripts were purified with a Monarch Spin RNA Isolation Kit. The concentrations of the resulting RNA stocks were determined on the spectrophotometer and adjusted to 2000 ng µL$^{-1}$ with nuclease-free water. The condensates were prepared by adding 1 µL Cy3-labeled RNA (2000 ng µL$^{-1}$) to 0.5 µL 400 mM $MgCl_2$, and 2.5 µL TE buffer. The resulting samples were subjected to the standard heating ramp. Afterward, they were gently vortexed, after which 2 µL was added to a 384-well glass-bottom plate (Corning) containing 28 µL 1× HEPES salt buffer. The condensates were left to sediment for at least 30 min before CLSM imaging. **For the incorporation of different RNAs into DNA-ACs,** 2.5 µL DNA-AC pre-mix A, 0.5 µL 400 mM $MgCl_2$, and 1 µL of the 2000 ng µL$^{-1}$ transcripts were placed in a 200 µL PCR tube and subjected to the standard heating ramp. Afterward, 2 µL y*-Atto 488 label (6.25 µM) and 2 µL x*-Atto 647N label (6.25 µM) were added and left to hybridize at RT for at least 30 min. The resulting dispersions were gently vortexed, after which 2 µL was added to a 384-well glass-bottom plate (Corning) containing 28 µL 1× HEPES salt buffer. The condensates were left to sediment for at least 30 min before CLSM imaging.

**Influence of RNA concentration on condensate morphology *(Figure 2a).*** Cy5-labeled AP3-RNA transcript (Transcription: 1 µL AP3 template (25 µM), 1 µL T7 promoter (25 µM), 2 µL 0.1 mM Aminoallyl-UTP-Cy5, 6.67 µL NTP Buffer mix, 1.33 µL T7 RNA Polymerase mix, and 8 µL nuclease-free water were gently homogenized and left to transcribe for 2 h at 37 °C) was purified using a Monarch Spin RNA Isolation Kit. The concentration of the resulting RNA stock was determined on the spectrophotometer, and 3 stocks of 3000, 1500, and 750 ng µL$^{-1}$ Cy5-labeled AP3-RNA in TE were prepared. In separate 200 µL PCR tubes, 1 µL of each RNA stock was added to 2.5 µL DNA-AC pre-mix A, and 0.5 µL 400 mM $MgCl_2$. The resulting samples had RNA concentrations of 750, 375 and 188 ng µL$^{-1}$, and were subjected to the standard heating ramp. Afterward, 2 µL y*-Cy3 label (6.25 µM) and 2 µL x*-Atto 488 label (6.25 µM) were added and left to hybridize at RT for at least 30 min. The resulting dispersions were gently vortexed, after which 2 µL was added to a 384-well glass-bottom plate (Corning) containing 28 µL 1× HEPES salt buffer. The condensates were left to sediment for at least 2 h before CLSM imaging.

**Fluorescence recovery after photobleaching (FRAP) *(Figure 2b-e).*** For FRAP experiments we used the DR-ACs prepared with 750 ng µL$^{-1}$ Cy5-labeled AP3-RNA, 144 ng µL$^{-1}$ poly($A_{20}$-x)$_n$, and 48 ng µL$^{-1}$ poly($T_{20}$-y)$_n$ from the previous section. The DR-ACs were first imaged using laser intensities below 2% at laser lines 488, 561, and 638 nm. Afterward, the ROI was bleached at 100% (for all lasers) twice, with an interval of 2:45 s, followed by low intensity imaging of the recovery with time intervals of 2 min 45 s. A second ROI was used to make a line profile of the DR-ACs before and after bleaching, using the Leica LAS X software (v4.3.0.24308).

**RNA organelle activity for selective cargo uptake and activation *(Figure 3a-d).*** For each DR-AC, a 200 µL PCR tube containing 2.5 µL DNA-AC pre-mix A, 0.5 µL 400 mM $MgCl_2$, and 1 µL unpurified transcript (Transcription: 1 µL AP3 template (25 µM) or Spinach template (25 µM), 1 µL T7 promoter (25 µM), 6.67 µL NTP Buffer mix, 1.33 µL T7 RNA Polymerase mix, and 10 µL nuclease-free water were gently homogenized and left to transcribe for 2 h at 37 °C), was subjected to the standard heating ramp. Afterward, 2 µL y*-Cy3 label (6.25 µM) was added to the AP3 DR-ACs and 2 µL y*-DY-647P1 label (6.25 µM) to the Spinach DR-

ACs, followed by a 30 min incubation time. The resulting dispersions were gently vortexed, after which 2 µL of each dispersion was added to a 384-well glass-bottom plate (Corning) containing 28 µL 1× HEPES salt buffer. The condensates were left to sediment for 2 h. To the AP3 DR-AC only, and AP3 DR-AC/Spinach DR-AC mix, 1 µL recombinant eGFP (33 µM) was added (final concentration: 1 µM eGFP), after which an immediate CLSM time-lapse was recorded. To the Spinach DR-AC only, and AP3 DR-AC/Spinach DR-AC mix, 1 µL DFHBI (150 µM) was added (final concentration: 5 µM DFHBI), after which an immediate CLSM time-lapse was recorded as well.

**Serum-mediated RNA degradation *(Figure 4a-e).*** A 200 µL PCR tube containing 2.5 µL DNA-AC pre-mix A, 0.5 µL 400 mM $MgCl_2$, and 1 µL purified Cy5-labeled AP3-RNA transcript (3000 ng $\mu L^{-1}$, protocol described under "*Influence of RNA concentration on condensate morphology*"), was subjected to the standard heating ramp. Afterward, 2 µL y*-Cy3 label (6.25 µM) and 2 µL x*-Atto 488 label (6.25 µM) were added and left to hybridize at RT for at least 30 min. The resulting dispersion was gently vortexed, after which 2 µL was added to a 384-well glass-bottom plate (Corning) containing 28 µL 1× HEPES salt buffer. The condensates were left to sediment for at least 30 min. To the well, 10 µL FBS was added (final concentration: 25% FBS) and quickly homogenized before a time-lapse was recorded on the CLSM.

**Serum-mediated protein release *(Figure 4f-h).*** A 200 µL PCR tube containing 1.5 µL DNA-AC pre-mix E, 0.5 µL 400 mM $MgCl_2$, 1.67 µL TE buffer and 0.33 µL AP3-RNA (purified, 2875 ng $\mu L^{-1}$, Biomers) was subjected to the standard heating ramp. Afterward, 2 µL y*-Cy3 label (6.25 µM, not shown in images), 2 µL x*-Atto 647N label (6.25 µM), and 1 µL recombinant eGFP (1 µM) were added and left to hybridize/diffuse at RT for at least 30 min. The resulting dispersion was gently vortexed, after which 2 µL was added to a 384-well glass-bottom plate (Corning) containing 28 µL 1× HEPES salt buffer. The condensates were left to sediment for at least 60 min. To the well, 10 µL FBS was added (final concentration: 25% FBS) and quickly homogenized before a time-lapse was recorded on the CLSM.

**Improved serum stability of RNA organelles by DNA shielding *(Figure 4i-k).*** Cy3-labeled AP3-RNA transcript (Transcription: 2 µL AP3 template (25 µM), 2 µL T7 promoter (25 µM), 6 µL 0.1 mM Aminoallyl-UTP-Cy3, 13.3 µL NTP Buffer mix, 2.67 µL T7 RNA Polymerase mix, and 14 µL nuclease-free water were gently homogenized and left to transcribe for 2 h at 37 °C) was purified using a Monarch Spin RNA Isolation Kit. The concentration of the resulting RNA stock was determined on the spectrophotometer and adjusted to 3000 ng $\mu L^{-1}$. The RNA condensates were prepared by adding 1 µL Cy3-labeled AP3-RNA (purified, 3000 ng $\mu L^{-1}$) to 0.5 µL 400 mM $MgCl_2$ and 2.5 µL TE buffer to a 200 µL PCR tube. The DR-ACs contained 1 µL Cy3-labeled AP3-RNA (purified, 3000 ng $\mu L^{-1}$), 0.5 µL 400 mM $MgCl_2$, and 2.5 µL DNA-AC pre-mix E in a 200 µL PCR tube. Both tubes were subjected to the standard heating ramp. Afterward, 2 µL y*-DY-647P1 label (6.25 µM) was added to the DR-ACs and 2 µL nuclease-free water was added to the RNA condensates. The samples were allowed to rest for 30 min, after which they were gently vortexed. 2 µL of each sample was added to the same well in a 384-well glass-bottom plate (Corning) that contained 26 µL 1× HEPES salt buffer and left to sediment for 2 h. To the well, 10 µL FBS was added (final concentration: 25% FBS) and quickly homogenized before a time-lapse was recorded on the CLSM.

**Synthesis and characterization of 2'F-modified RNA *(Figure S6).*** The 2'F-modified RNA was generated from an AP3 dsDNA template that was prepared by PCR. Reactions were performed using Phusion High-Fidelity DNA Polymerase with 1 µM AP3 forward and reverse primers (Table S2). The thermal cycling program was as follows: initial denaturation at 95 °C for 2 min, followed by cycles of 95 °C for 30 s, 55 °C for 30 s, and 72 °C for 20 s with a final extension step at 72 °C for 2 min. PCR products were purified using the GeneJET PCR purification kit. AP3-2'OH and AP3-2'F were generated by transcription from the purified dsDNA (0.7 µM final concentration). The transcription mix contained Y639F T7 RNA polymerase (1x) in T7 transcription buffer (80 mM HEPES pH 7.5, 25 mM $MgCl_2$, 2 mM spermidine-HCl) with ATP, GTP, and either CTP/UTP or 2′F-dCTP/2′F-dUTP (2.5 mM each). For Cy5 and 2'F-containing AP3 transcripts, a 95:5 (%) mixture of 2′F-dUTP:Cy5-UTP (2.5 mM total) was used. Reactions were supplemented with 12.5 mM DTT, 0.005 U/µL inorganic pyrophosphatase, and 0.05 mg/mL BSA, brought to volume with nuclease-free water, and incubated overnight at 37 °C. RNA products were purified using the RNA Clean & Concentrator kit.

The binding affinity of unmodified AP3 (AP3-2'OH) and 2'-F modified AP3 (AP3-2'F) was measured using BLI. His-tagged eGFP (10 ng/µL) in BLI buffer (40 mM HEPES, 125 mM KCl, 5 mM $MgCl_2$ supplemented with 0.1 mg/mL BSA and 0.02% Tween20) was immobilized onto OCTET Ni-NTA Biosensors. Serial dilutions of the aptamers, previously folded (95 °C for 2 min, 65 °C for 5 min, 37 °C for 5 min, and held at 21 °C), were prepared in BLI buffer. Baseline signals were recorded prior to each binding event (including association, dissociation, and regeneration steps). The eGFP-coated sensor was dipped into the aptamer solution for 400 s (association), followed by dipping into buffer alone for 1200 s (dissociation). Regeneration was performed over three cycles, each consisting of a 5 s dip in glycine solution (10 mM at pH 1.4) followed by a 5 s dip in BLI buffer. Sensorgrams were baseline-aligned for further analysis.

For serum stability evaluation of the two AP3 versions, refolded aptamers were incubated at 20 µM in 40 mM HEPES, 125 mM KCl, and 5 mM $MgCl_2$ with 25% FBS at RT. Samples were collected at 0, 4 min, and 1, 4, 24, 48 and 120 h. The samples were diluted 1:10 in nuclease-free water, and 3 pmol was analyzed by 2% agarose gel electrophoresis.

**Serum-stable RNA condensates via 2'F-modified RNA *(Figure S7)*.** In 200 µL PCR tubes, 1 µL Cy3-labeled AP3-RNA (purified, 3000 ng µL$^{-1}$, synthesis above) or 1 µL Cy5-2'F-modified AP3-RNA (purified, 3000 ng µL$^{-1}$, synthesis above) was added to 0.5 µL 400 mM $MgCl_2$ and 2.5 µL TE buffer. Both tubes were subjected to the standard heating ramp. The resulting dispersions were gently vortexed. 2 µL of each condensate was added to the same well in a 384-well glass-bottom plate (Corning) that contained 26 µL 1× HEPES salt buffer and left to sediment for 2 h. To the well, 10 µL FBS was added (final concentration: 25% FBS) and quickly homogenized before a time-lapse was recorded on the CLSM.

**Serum-stable DR-ACs via 2'F-modified RNA *(Figure 4l-n and S8)*.** The unmodified DR-ACs were prepared by adding 1.25 µL DNA-AC pre-mix E, 0.5 µL 400 mM $MgCl_2$, 1.75 µL TE buffer, and 0.5 µL purified AP3-RNA (purified, 2875 ng µL$^{-1}$, Biomers) to a 200 µL PCR tube. The 2'F-modified DR-ACs contained 1.25 µL DNA-AC pre-mix E, 0.5 µL 400 mM $MgCl_2$, 1.65 µL TE buffer, and 0.6 µL 2'F-modified AP3-RNA (purified, 1340 ng µL$^{-1}$, prepared as described above). Both tubes were subjected to the standard heating ramp. Afterward, 1 µL recombinant eGFP (1 µM) was added to each sample, 2 µL y*-Cy3 label (6.25 µM) was added to the unmodified DR-ACs, and 2 µL y*-DY-647P1 was added to the 2'F-modified DR-ACs. The samples were allowed to rest for 30 min, after which they were gently vortexed. 2 µL of each sample was added to the same well in a 384-well glass-bottom plate (Corning) that contained 26 µL 1× HEPES salt buffer and left to sediment for 2 h. To the well, 10 µL FBS was added (final concentration: 25% FBS) and quickly homogenized before a time-lapse was recorded on the CLSM. A negative control without FBS was included.

**Sender DNA-AC-activated gene expression *(Figure 5a-d)*.** The Sender DNA-ACs were prepared by adding 4 µL DNA-AC pre-mix A, 1 µL 400 mM $MgCl_2$, and 4 µL x*-P*-A*-x* (10 µM) to a 200 µL PCR tube. The tube was subjected to the standard heating ramp. Afterward, 4 µL y*-Cy3 label (6.25 µM) was added and left to hybridize at RT for at least 30 min. In a 384-well clear-bottom polystyrene plate with a non-binding surface, two samples were prepared in triplicate. Sample 1: From the PURExpress TX-TL kit; 4.9 µL Solution A and 3.65 µL Solution B. 1 µL of the freshly prepared Sender DNA-ACs, 2 µL gated eGFP mRNA template plasmid (125 ng µL$^{-1}$), 2 µL T7 promoter (5 µM), and 6.45 µL nuclease-free water. Sample 2 (negative control): From the PURExpress TX-TL kit; 4.9 µL Solution A and 3.65 µL Solution B. 1 µL of the freshly prepared Sender DNA-ACs, 2 µL gated eGFP mRNA template plasmid (125 ng µL$^{-1}$), and 8.45 µL nuclease-free water. The samples were overlaid with 10 µL hexadecane and measured in the plate reader at 37 °C.

**Multi-cell-like communication via Sender-Receiver *(Figure 5e,f)*.** The Sender DNA-AC was prepared as described above. The Receiver DR-AC was prepared by adding 2 µL AP3-RNA (purified, 2875 ng µL$^{-1}$, Biomers), 1 µL 400 mM $MgCl_2$, and 5 µL DNA-AC pre-mix A to a 200 µL PCR tube. The tube was subjected to the standard heating ramp. Afterward, 4 µL y*-Cy3 label (6.25 µM) was added and left to hybridize at RT for at least 30 min. Next, 3 separate samples were assembled in triplicate in a 384-well glass-bottom plate (Corning). Sample A: From the PURExpress TX-TL kit; 4.9 µL Solution A and 3.65 µL Solution B. 1 µL of the freshly prepared Sender DNA-ACs, 1 µL of the freshly prepared Receiver DR-ACs, 2 µL gated eGFP mRNA template plasmid (125 ng µL$^{-1}$), and 5.45 µL nuclease-free water. Sample B (negative control 1): From the PURExpress TX-TL kit; 4.9 µL Solution A and 3.65 µL Solution B. 1 µL of the freshly prepared Sender DNA-ACs, 1 µL of the freshly prepared Receiver DR-ACs, 2 µL gated eGFP mRNA template plasmid (125 ng µL$^{-1}$), and 7.45 µL nuclease-free water. Sample C (negative control 2): From the PURExpress TX-TL kit; 4.9 µL Solution A and 3.65 µL Solution B. 1 µL of the freshly prepared Receiver DR-ACs, 2 µL gated eGFP mRNA template plasmid (125 ng µL$^{-1}$), and 6.45 µL nuclease-free water. The ACs were allowed to sediment for 30 min, after which the $t = 0$ images were recorded on the CLSM. Afterward, 2 µL DY-647P1-labeled T7 promoter

(5 µM) was added to the A and C samples, after which the well plate was incubated at 37 °C for 90 min, which was directly followed by a second round of CLSM imaging.

**Kinetics of multi-cell-like communication via Sender-Receiver *(Figure 5g)*.** The Sender DNA-ACs were prepared by adding 6 µL DNA-AC pre-mix A, 1.5 µL 400 mM $MgCl_2$, and 6 µL x*-P*-A*-x* (10 µM) to a 200 µL PCR tube. The Receiver DR-AC was prepared by adding 3 µL AP3-RNA (purified, 2875 ng $µL^{-1}$, Biomers), 1.5 µL 400 mM $MgCl_2$, and 7.5 µL DNA-AC pre-mix A to a 200 µL PCR tube. The tubes were subjected to the standard heating ramp. Afterward, 6 µL nuclease-free water was added to both tubes. Four identical samples were prepared in 200 µL PCR tubes: From the PURExpress TX-TL kit; 18.4 µL Solution A and 13.7 µL Solution B. 3.75 µL of the freshly prepared Sender DNA-ACs, 3.75 µL of the freshly prepared Receiver DR-ACs, 7.5 µL gated eGFP mRNA template plasmid (125 ng $µL^{-1}$), and 27.9 µL nuclease-free water. An aliquot of 2.5 µL was taken of each sample, which was then diluted with 97.5 µL 1× HEPES salt buffer, and directly measured with flow cytometry. Afterward, 0.375 µL DY-647P1-labeled promoter (100 µM) was added to two of the samples. To the two remaining samples, 0.375 µL nuclease-free water was added, turning them into the negative controls. All samples were placed in a ThermoMixer at 37 °C for 95 min. During this time, the samples were gently homogenized every 5 min by gentle pipetting, and an aliquot of 2.5 µL was taken of each sample, which was subsequently diluted with 97.5 µL 1× HEPES salt buffer, and directly measured with flow cytometry. For flow cytometry measurements approximately 500 ACs were measured per second with a cut-off at 10,000 events in the earlier-identified AC gated region. By plotting the Sender DNA-AC response (DY-647P1; RL637nm 667/30-A) against the Receiver DR-AC response (eGFP; BL488nm 525/45-A), the particles could be identified. The region for the Receiver DR-ACs was identically gated for each time point, providing a mean eGFP fluorescence intensity of the Receiver DR-AC particles over time.

**Integrated signal processing in single DR-ACs *(Figure 5h,i)*.** The self-actuating DR-AC was prepared by adding 2 µL AP3-RNA (purified, 2875 ng $µL^{-1}$, Biomers), 1 µL x*-P*-A*-x* (10 µM), 1.2 µL 400 mM $MgCl_2$, and 5 µL DNA-AC pre-mix A to a 200 µL PCR tube. The tube was subjected to the standard heating ramp. Afterward, 4 µL y*-Cy3 label (6.25 µM) was added and left to hybridize at RT for at least 30 min. Next, 2 separate samples were assembled in triplicate in a 384-well glass-bottom plate (Corning). Sample A: From the PURExpress TX-TL kit; 4.9 µL Solution A and 3.65 µL Solution B. 1 µL of the freshly prepared self-actuating DR-ACs, 2 µL gated eGFP mRNA template plasmid (125 ng $µL^{-1}$), and 6.45 µL nuclease-free water. Sample B (negative control 1): From the PURExpress TX-TL kit; 4.9 µL Solution A and 3.65 µL Solution B. 1 µL of the freshly prepared self-actuating DR-ACs, 2 µL gated eGFP mRNA template plasmid (125 ng $µL^{-1}$), and 8.45 µL nuclease-free water. The DR-ACs were allowed to sediment for 30 min, after which the $t$ = 0 images were recorded on the CLSM. Afterward, 2 µL DY-647P1-labeled T7 promoter (5 µM) was added to sample A, after which samples were incubated at 37 °C for 90 min, followed by CLSM imaging.

### 4. Supplementary Tables

**Table S1: DNA sequence for gated eGFP mRNA oligonucleotide sequences** as synthesized and inserted in the plasmid via GenScript. The essential regions are marked by color. T7 promoter (Yellow), Activator Binding Site (Magenta, A*), Ribosome Binding Site (Blue), and eGFP gene (Green).

| nt Number | Sequence (5'→ 3') |
|---|---|
| 1 | ATGCATGTCT TCCGCGGTAA TACGACTCAC TATAGGGTCT TATCTTATCT ATCTCGTTTA |
| 61 | TCCCTGCATA CAGAAACAGA GGAGATATGC AATGATAAAC GAGAACCTGG CGGCAGCGCA |
| 121 | AAAGATGGTG AGCAAGGGCG AGGAGCTGTT CACCGGGGTG GTGCCCATCC TGGTCGAGCT |
| 181 | GGACGGCGAC GTAAACGGCC ACAAGTTCAG CGTGTCCGGC GAGGGCGAGG |
| 241 | GCGATGCCAC |
| 301 | CTACGGCAAG CTGACCCTGA AGTTCATCTG CACCACCGGC AAGCTGCCCG TGCCCTGGCC |
| 361 | CACCCTCGTG ACCACCCTGA CCTACGGCGT GCAGTGCTTC AGCCGCTACC CCGACCACAT |
| 421 | GAAGCAGCAC GACTTCTTCA AGTCCGCCAT GCCCGAAGGC TACGTCCAGG AGCGCACCAT |
| 481 | CTTCTTCAAG GACGACGGCA ACTACAAGAC CCGCGCCGAG GTGAAGTTCG AGGGCGACAC |
| 541 | CCTGGTGAAC CGCATCGAGC TGAAGGGCAT CGACTTCAAG GAGGACGGCA ACATCCTGGG |
| 601 | GCACAAGCTG GAGTACAACT ACAACAGCCA CAACGTCTAT ATCATGGCCG ACAAGCAGAA |
| 661 | GAACGGCATC AAGGTGAACT TCAAGATCCG CCACAACATC GAGGACGGCA GCGTGCAGCT |
| 721 | CGCCGACCAC TACCAGCAGA ACACCCCCAT CGGCGACGGC CCCGTGCTGC TGCCCGACAA |
| 781 | CCACTACCTG AGCACCCAGT CCGCCCTGAG CAAAGACCCC AACGAGAAGC GCGATCACAT |
| 841 | GGTCCTGCTG GAGTTCGTGA CCGCCGCCGG GATCACTCTC GGCATGGACG AGCTGTACAA |
| | GTAGAGTTCT GCACGATATC GTCTTATGCA TGTCTTCCGC GGATACAC |

**Table S2: Oligonucleotide sequences** as purchased from Biomers or IDT, with their name, sequence, purification method and modifications.

| Name | Sequence (5'→ 3') | Purification | Modification |
|---|---|---|---|
| **Template ($A_{20}$-x) (DNA)** | ATC TAT CCT AAT TTT TTT TTT TTT TTT TTT TGA ACC CGT AT | HPLC | 5' phosphate |
| **x ligation (DNA)** | TTA GGA TAG ATA TAC GGG TTC | Cartridge | none |
| **Primer-x (DNA)** | TTA GGA TAG ATA TAC GGG T*T *C | Cartridge | *=PTO |
| **Template ($T_{20}$-y) (DNA)** | ATC CTC TAA AAT CAA AAA AAA AAA AAA AAA AAG TAA AAC CAC ACG | HPLC | 5' phosphate |
| **y ligation (DNA)** | TTT TAG AGG ATC GTG TGG TTT T | Cartridge | none |
| **Primer-y (DNA)** | TTT TAG AGG ATC GTG TGG TT* T*T | Cartridge | *=PTO |
| **x*-Atto 488 (DNA)** | TGA ACC CGT ATA TCT ATC CTA A | HPLC | 5' Atto 488 |
| **x*-Atto 647N (DNA)** | TGA ACC CGT ATA TCT ATC CTA A | HPLC | 5' Atto 647N |
| **y*-Atto 488 (DNA)** | AAA ACC ACA CGA TCC TCT A | HPLC | 5' Atto 488 |
| **y*-Cy3 (DNA)** | AAA ACC ACA CGA TCC TCT A | HPLC | Cyanine 3 |
| **y*-DY-647P1 (DNA)** | AAA ACC ACA CGA TCC TCT A | HPLC | 5' DY-647P1 |
| **T7 promoter (DNA)** | TAA TAC GAC TCA CTA TA | Cartridge | none |
| **DY-647P1-labeled T7 promoter (DNA)** | TAA TAC GAC TCA CTA TA | HPLC | 5' DY-647P1 |
| **AP3 (RNA)** | GGG AGC UUC UGG ACU GCG AUG GGA GCA CGA AAC GUC GUG GCG CAA UUG GGU GGG GAA AGU CCU UAA AAG AGG GCC ACC ACA GAA GCU | HPLC | none |
| **[A] *Random* Template (DNA)** | ATT GTG CGA TGT CCT CGA TCT CTA CCT CCA TCC CTA TAG TGA GTC GTA TTA GCG AGT ATA GGG | HPLC | none |
| **[B] *Random* Template (DNA)** | CCA GGC ACT GAC ATT CCA CGA AGG CGC CAA TAT ACC CTT TCC CTA TAG TGA GTC GTA TTA | Cartridge | none |
| **[C] x*-P*-A*-x* (DNA)** | GAA CCC GTA TAT CTA TCC TAA ATC TTA TCT ATC TCG TTT ATC CCT GCC CTA TAG TGA GTC GTA TTA GGG GGC TAA TAC TAT AGG GGA ACC CGT ATA TCT ATC CTA A | HPLC | none |
| **[D] *Spinach* Template (DNA)** | GGA GCT CAC ACT CTA CTC AAC AGT AGC GAA CTA CTG GAC CCG TCC TTC ACC CTA TAG TGA GTC GTA TTA GCG AGT ATA GGG | HPLC | none |
| **[E] *Random* Template (DNA)** | ACT GCA CGT CCA GGC ACT GAC ATT CCA CGA AGG CGC CCA ATA TAC CAG TAT CGA TCG GAC GTG AGC GTA CGT GAG CGT CCC TAT AGT GAG TCG TAT TA | HPLC | none |
| **[F] *AP3* Template (DNA)** | AGC TTC TGT GGT GGC CCT CTT TTA AGG ACT TTC CCC ACC CAA TTG CGC CAC GAC GTT TCG TGC TCC CAT CGC AGT CCA GAA GCT CCC TAT AGT GAG TCG TAT TA | HPLC | none |
| **[G] *Random* Template (DNA)** | ACT ACA GCC GCA CAA GAA ACC CAG AAC ATC AGA CCA ACA CAA GAA ACC CAA CCC AGA ACA TCA TTG CTC CTC TTA CGT CAT TAT TCA TCA GTA CTA CCC GTC CTT CAC CCT ATA GTG AGT CGT ATT AGC GAG TAT AGG G | HPLC | none |

| | | | |
|---|---|---|---|
| **[H] Forward Primer (DNA)** | TAA TAC GAC TCA CTA TAG GGA GTC TAC CTC CCG CCA TAA AAA | Cartridge | none |
| **[H] Reverse Primer (DNA)** | TTA TGT TTC AGG TTC AGG GGG A | Cartridge | none |
| **AP3 Primer (Forward)** | AG CTT CTG TGG TGG CCC TCT | Cartridge | none |
| **AP3 Primer (Reverse)** | TAA TAC GAC TCA CTA TAG GGA GCT TCT GGA CTG CGA TGG GAG CAC | Cartridge | none |

### 5. Supplementary Figures

a

| Name | Sequence |
|---|---|
| $A_{20}$-x | AAA AAA AAA AAA AAA AAA AA TTA GGA TAG ATA TAC GGG TTC |
| $T_{20}$-y | TT TTT TTT TTT TTT TTT TTT GA TTT TAG AGG ATC GTG TGG TTT T AC |
| AP3 | GGG AGC UUC UGG ACU GCG AUG GGA GCA CGA AAC GUC GUG GCG CAA UUG GGU GGG GAA AGU CCU UAA AAG AGG GCC ACC ACA GAA GCU |

b

c

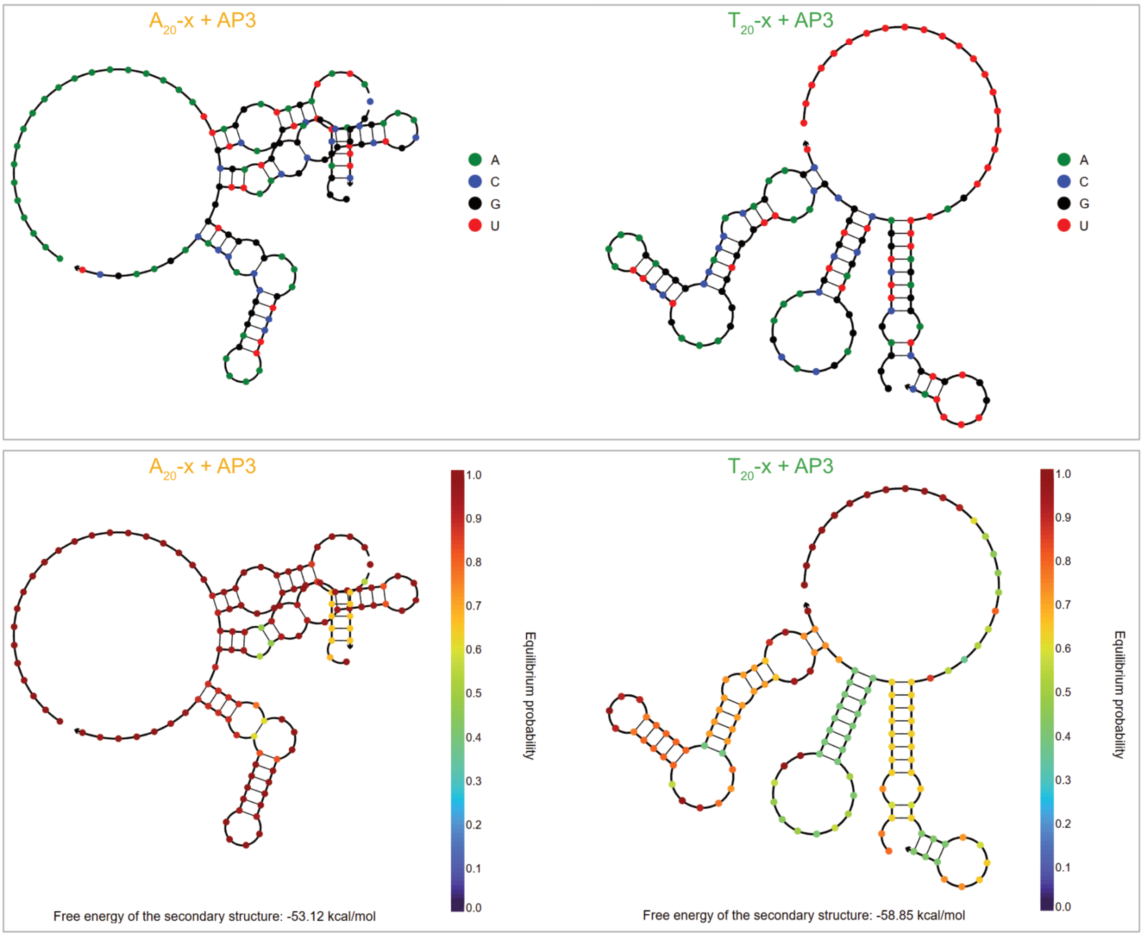

d

| Name | Concentration total (M) | Concentration complexed with AP3 (M) | Percentage complexed with AP3 (%) |
|---|---|---|---|
| $A_{20}$-x | 9.00E-06 | 2.66E-09 | 0.105 |
| $T_{20}$-y | 3.00E-06 | 9.43E-09 | 0.314 |
| AP3 | 1.70E-05 | — | — |

**Figure S1. NUPACK simulations showing minimal specific interactions between AP3 and $A_{20}$-x and $T_{20}$-y sequences.** a) Table with sequences of AP3-RNA (AP3), and single repeating units of poly($A_{20}$-x)$_n$ = $A_{20}$-x and poly($T_{20}$-y)$_n$ = $T_{20}$-y. b) NUPACK-simulated MFE RNA proxy structures $A_{20}$-x/AP3 (left) and $T_{20}$-y/AP3 (right) complexes represented in nucleotide identity, and in equilibrium probability (c). d) Table quantifying simulated $A_{20}$-x/AP3 and $T_{20}$-y/AP3 complex formations. A simulation with 9.00 µM $A_{20}$-x and 1.70E-05 M AP3 predicts 2.66 E-09 M $A_{20}$-x to be complexed with AP3, which is 0.105% of the $A_{20}$-x present. A simulation with 3.00E-06 M $T_{20}$-y and 1.70E-05 M AP3 predicts 9.43 E-09 M $T_{20}$-y to be complexed with AP3, which is 0.314% of the $T_{20}$-y present. NUPACK simulations were performed with oligonucleotide concentrations presented in the table in panel (d), model parameter: rna06, at 10 °C.

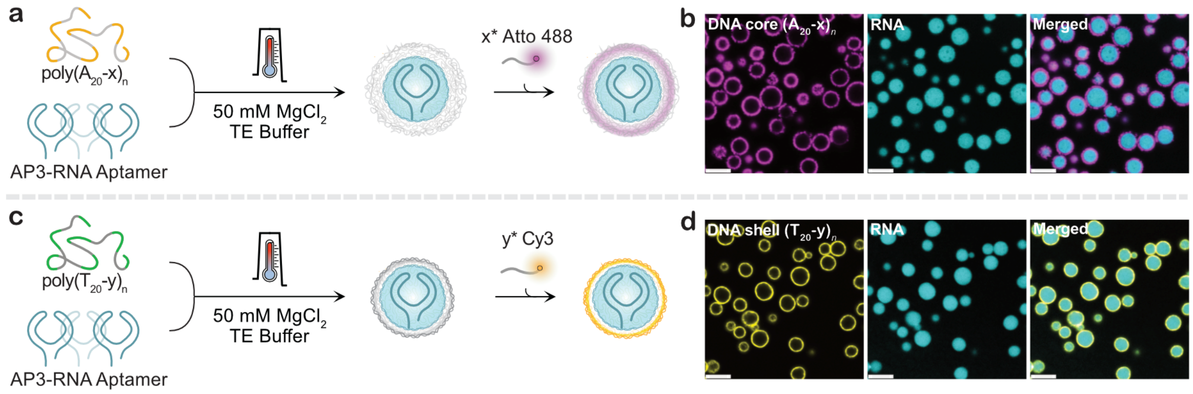

**Figure S2. Formation of DR-ACs from RNA and either poly($A_{20}$-x)$_n$ or poly($T_{20}$-y)$_n$.** a) Schematic illustration of DR-AC assembly from Cy5-labeled AP3-RNA (unpurified transcript) and 144 ng µL$^{-1}$ poly($A_{20}$-x)$_n$, subsequently labeled with x*-Atto 488; corresponding CLSM images are shown in (b). c) Schematic illustration of DR-AC assembly from Cy5-labeled AP3-RNA (unpurified transcript) and 48 ng µL$^{-1}$ poly($T_{20}$-y)$_n$, subsequently labeled with y*-Cy3; corresponding CLSM images are shown in (d). All CLSM images were acquired after at least a 30 min incubation in 40 mM HEPES, 125 mM KCl and 5 mM $MgCl_2$ at RT. Scale bar: 5 µm.

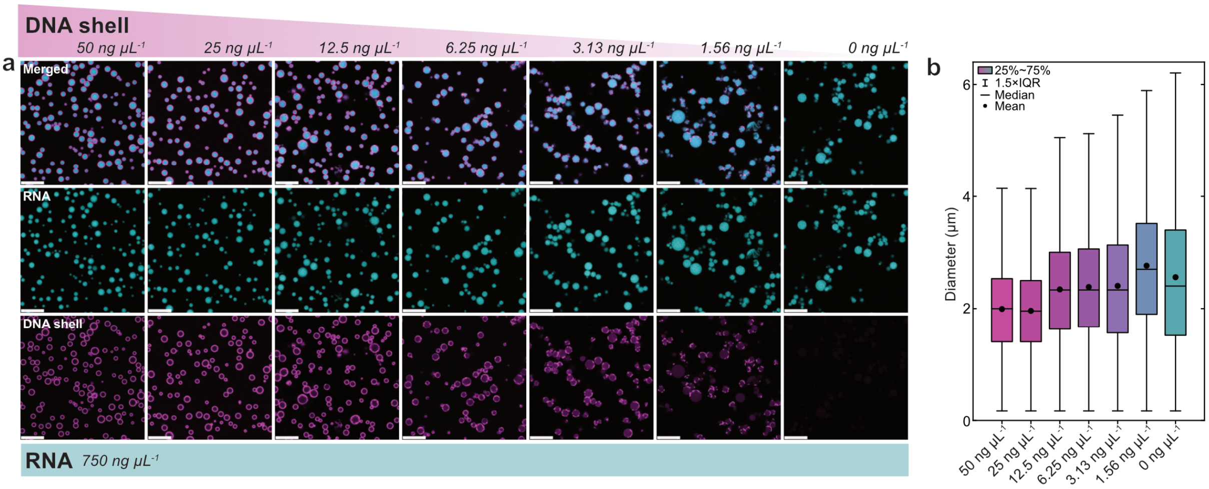

**Figure S3. Effect of DNA shell concentration on RNA condensate size distribution and homogeneity.** a) CLSM images of AP3-RNA–poly($T_{20}$-y)$_n$ DR-ACs prepared at a constant concentration (750 ng µL$^{-1}$) of Cy5-labeled AP3-RNA and varying poly($T_{20}$-y)$_n$ concentrations (50, 25, 12.5, 6.25, 3.13, 1.56 or 0 ng µL$^{-1}$). Decreasing the DNA shell concentration results in increased RNA condensate coalescence in the heating ramp and patchy poly($T_{20}$-y)$_n$ shell structures. Reduced poly($T_{20}$-y)$_n$ concentrations lead to increased particle diameters and size distributions. Labeling: y*-Atto 488. All CLSM images were acquired after at least 30 min incubation in 40 mM HEPES, 125 mM KCl and 5 mM $MgCl_2$ at RT. Scale bar: 10 µm. b) Corresponding AP3-RNA condensate diameters calculated from 8 respective images per condition (n = 1781 – 5707).

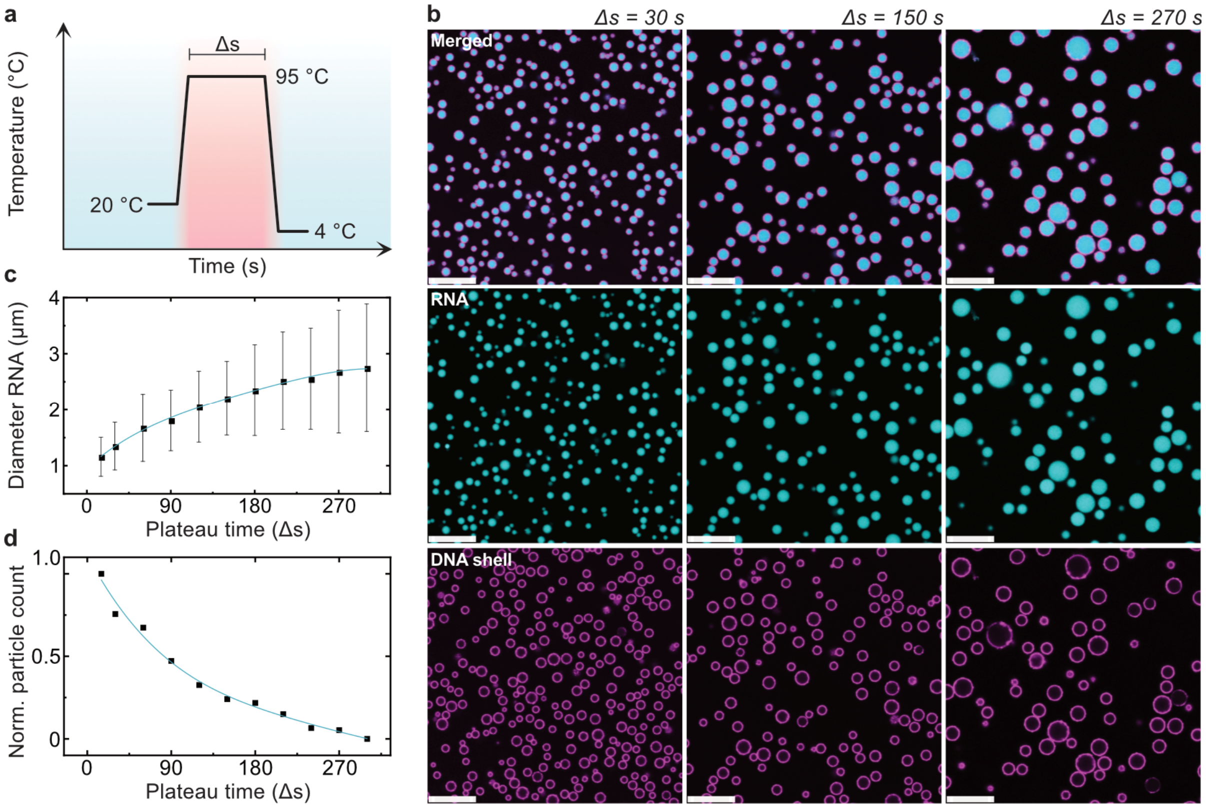

**Figure S4. Influence of temperature ramp plateau time on RNA condensate size and size distribution in DR-ACs.** a) Schematic illustration of temperature ramp used to synthesize DR-ACs. The temperature is rapidly increased (8 °C $s^{-1}$) from 20 °C to 95 °C, held for varying plateau times (Δs), and then rapidly cooled to 4 °C (6 °C $s^{-1}$). b) Representative CLSM images of AP3-RNA–poly($T_{20}$-y)$_n$ DR-ACs (750 ng µ$L^{-1}$ Cy5-labeled AP3-RNA and 25 ng µ$L^{-1}$ poly($T_{20}$-y)$_n$) synthesized with plateau times of 30, 150 and 270 s (Δs) and labeled with y*-Atto 488. c) Average diameter and size distribution of the RNA condensates in AP3-RNA–poly($T_{20}$-y)$_n$ DR-ACs synthesized with Δs = 15, 30, 60, 90, 120, 150, 180, 210, 240, 270 and 300 s. RNA condensate diameters and size distributions increase with longer plateau times. d) Correlation between plateau time (Δs) and normalized counts of RNA condensates per image frame at the same magnification. After the temperature ramp, equal volumes of product were left to incubate for 2 h in 40 mM HEPES, 125 mM KCl and 5 mM $MgCl_2$ at RT. For each sample, 3 CLSM images (34,047 µ$m^2$ each) were captured, and all RNA condensates were counted with counts ranging from n = 12,850 (Δs = 15 s) to n = 983 (Δs = 300 s). Scale bars: 10 µm.

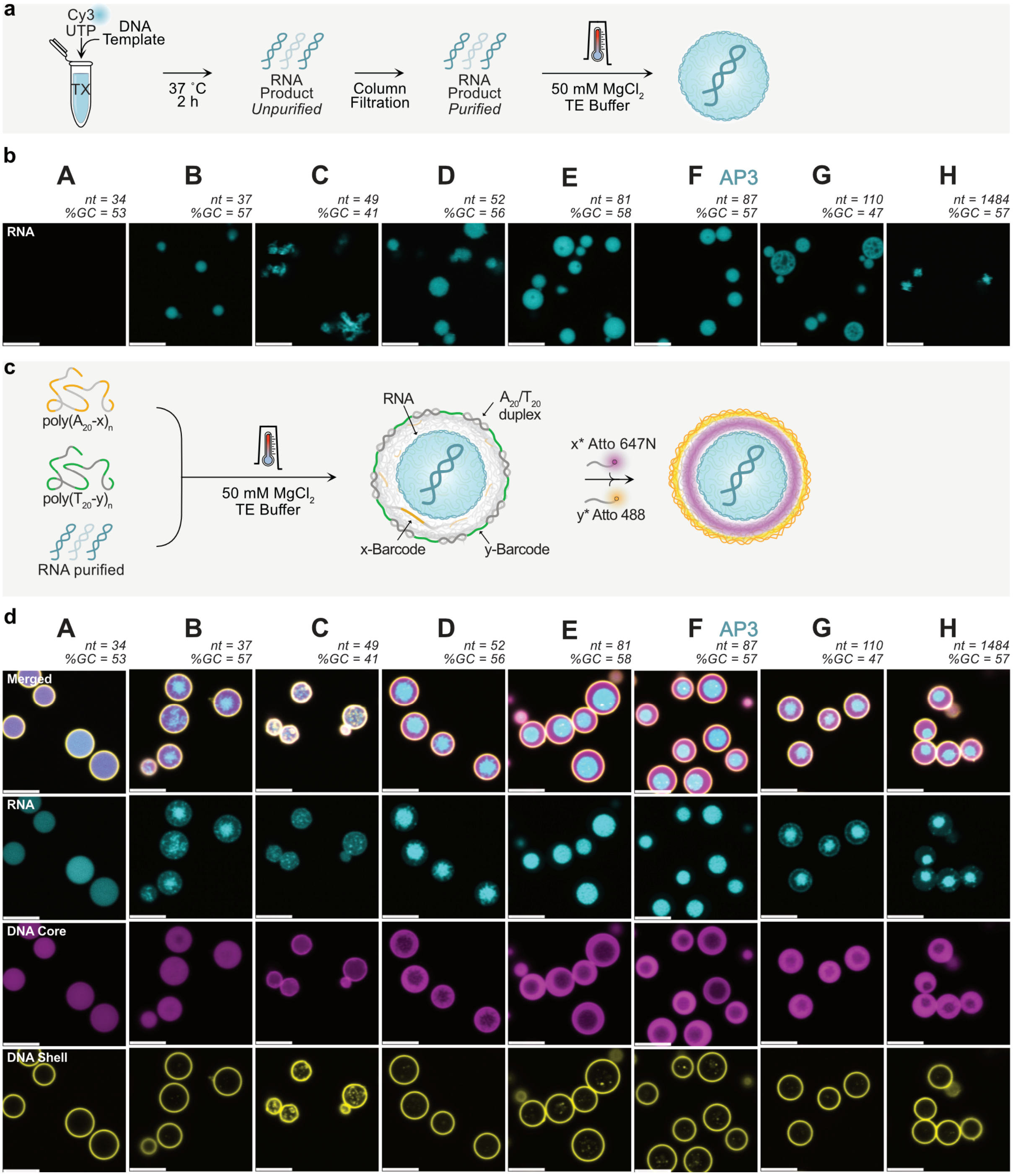

**Figure S5. DR-ACs formed with varying RNA sequences.** a) Schematic illustration of RNA condensate formation from purified Cy3-labeled RNA strands with varying lengths (*nt*) and GC content (*%GC*). b) CLSM images of RNA condensates generated from DNA templates A-H (sequences in Table S2, final RNA concentrations: 500 ng µL$^{-1}$). The resulting RNA condensates vary in quality (packing density and roundness), whereas RNA from template A yields no detectable condensates. c) Schematic illustration of DR-AC assembly from purified RNA products generated in panel a (500 ng µL$^{-1}$), 144 ng µL$^{-1}$ poly($A_{20}$-x)$_n$ post-labeled with an x*-Atto 647N label, and 48 ng µL$^{-1}$ poly($T_{20}$-y)$_n$ post-labeled with y*-Atto 488 label. d) CLSM images of corresponding DR-ACs. All DR-ACs contain RNA, albeit not always as a single, solid organelle core. The quantity and morphology of RNA differ per DR-AC type. All CLSM images were acquired after at least 30 min incubation in 40 mM HEPES, 125 mM KCl and 5 mM $MgCl_2$ at RT. Scale bars: 5 µm.

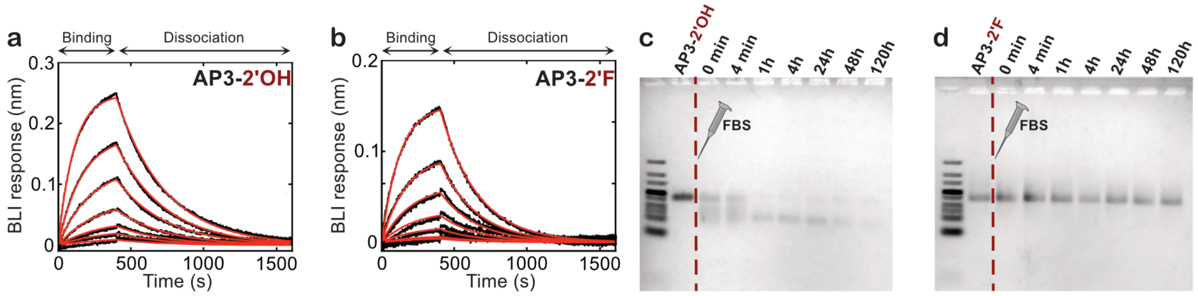

**Figure S6. Enhanced serum stability of 2'F-modified AP3-RNA with maintained eGFP binding.** Biolayer interferometry measurements of binding to eGFP-His (immobilized) for a) AP3-2'OH and b) AP3-2'F in solution with 400 s association and 1200 s dissociation times and aptamer concentrations ranging from 150 to 0 nM in a ½ serial dilution. Corresponding $K_D$ values are 69.4 nM for AP3-2'OH and 139.8 nM for AP3-2'F. Agarose gel electrophoresis (2 wt%) showing time-course degradation of AP3-RNA in 25% FBS for c) unmodified AP3-RNA (AP3-2'OH), and d) 2'F-modified AP3-RNA (AP3-2'F).

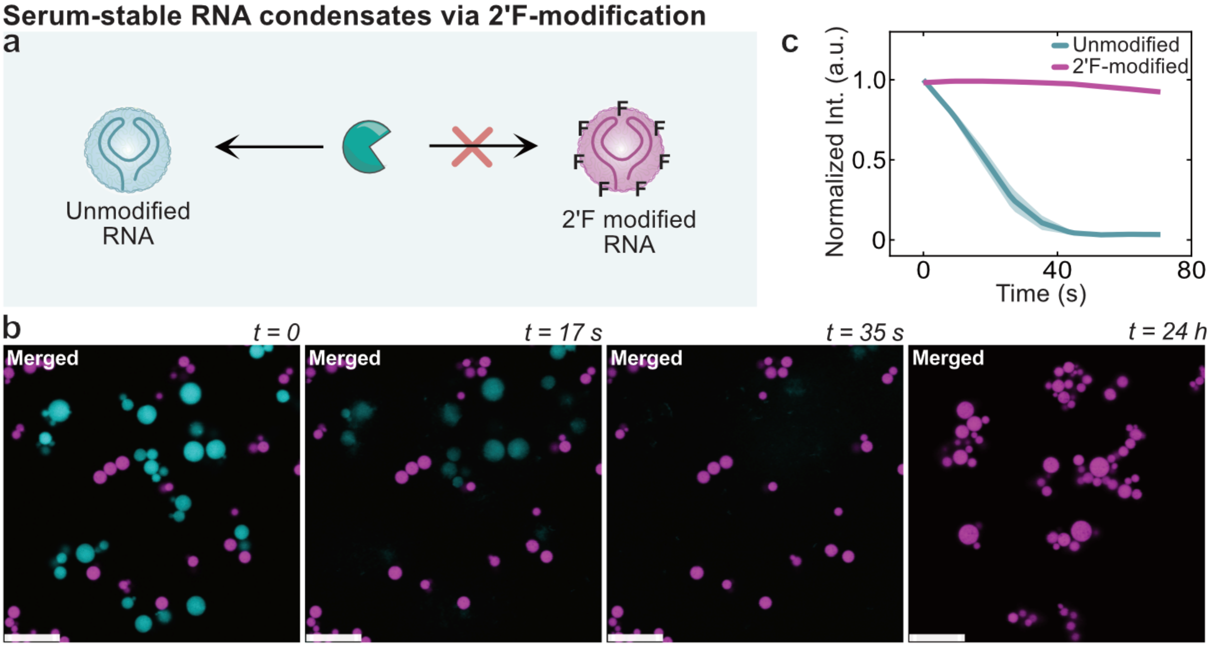

**Figure S7. Increased serum stability of RNA condensates with 2'F-modified RNA.** a) Schematic illustration showing that RNase digests unmodified RNA condensates but leaves 2'F-modified RNA condensates intact. b) Real-time CLSM images following addition of 25% FBS to a 1:1 mixture of unmodified RNA condensates (cyan) and 2'F-modified RNA condensates (magenta) over time (image at *t* = 24 h was taken from a different region in the same well). c) Corresponding fluorescence intensity plots of Cy3 (unmodified RNA condensates) and Cy5 (2'F-modified RNA condensates) for n = 8 condensates each (shaded areas represent the standard deviation). The fluorescent signal of the unmodified RNA condensates fully disappears within 40 s, whereas the 2'F-modified RNA condensates remain intact. (Conditions: unmodified RNA condensates: 750 ng $\mu L^{-1}$ AP3-Cy3 RNA, 2'F-modified condensates: 750 ng $\mu L^{-1}$ AP3-Cy5-2'F RNA, in 25% FBS). 40 mM HEPES, 125 mM KCl and 5 mM $MgCl_2$. Scale bar: 10 µm.

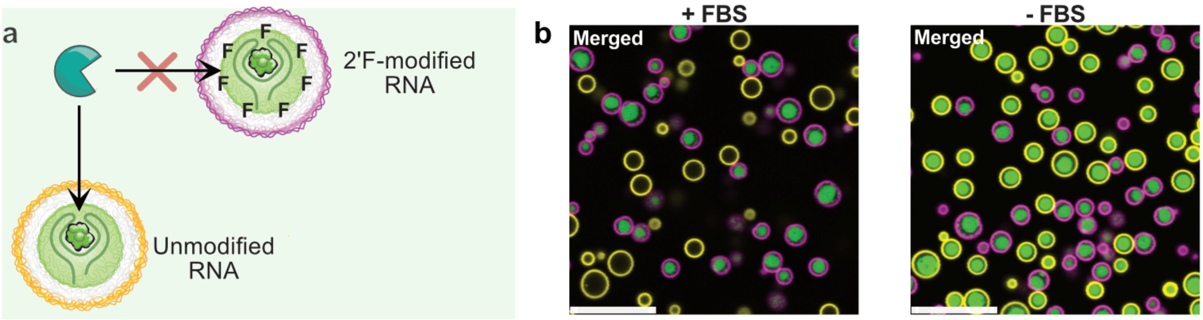

**Figure S8. Serum-stable DR-ACs via 2'F-modified RNA and negative control.** a) Schematic illustration of the stabilization effect of incorporated 2'F-modifications in the RNA organelle in DR-ACs against FBS-mediated degradation. b) CLSM images of a 1:1 mixture of unmodified (yellow shell) and 2'F-modified (magenta shell) AP3 DR-ACs 24 h after addition of 25% FBS (left) or without FBS (right), incubated at RT and measured at the same laser settings. (Compositions: unmodified DR-AC: 359 ng $\mu L^{-1}$ unmodified AP3-RNA, 72 ng $\mu L^{-1}$ poly($A_{20}$-x)$_n$, and 18 ng $\mu L^{-1}$ poly($T_{20}$-y)$_n$ (20:4:1 ratio). 2'F-modified DR-AC: 201 ng $\mu L^{-1}$ 2'F-modified AP3-RNA, 72 ng $\mu L^{-1}$ poly($A_{20}$-x)$_n$, and 18 ng $\mu L^{-1}$ poly($T_{20}$-y)$_n$ (11:4:1 ratio), (labeling: DNA shell unmodified DR-ACs; y*-Cy3 label, DNA shell 2'F-modified DR-ACs; y*-DY-647P1 label, RNA organelles; 7 nM eGFP) in 25% FBS (positive control only)). Buffer: 40 mM HEPES, 125 mM KCl and 5 mM $MgCl_2$. Scale bars: 10 µm.
